# Evolution towards small colony variants of pandemic multidrug resistant ST131 *Escherichia coli* isolates from a 10-year wound infection

**DOI:** 10.1101/2022.05.07.487787

**Authors:** Ali Dadvar, Shaiqa Labiba, Fengyang Li, Sulman Shafeeq, Marcus Ahl, Petra Lüthje, Natalia O. Dranenko, Janja Trcek, Anna Marzec-Grządziel, Sofya K. Garushyants, Hanna Nagel, Irfan Ahmad, Armin O. Schmitt, Analucia Diaz Lostao, Volkan Özenci, Måns Ullberg, Mikhail S. Gelfand, Börje Åkerlund, Ute Römling

## Abstract

Chronic wounds are difficult to treat because underlying medical conditions can impair the mechanical and physiological first-line innate immune defenses, leading to persistent microbial infections. We report here the isolation, molecular and phenotypic characterization of seven *E. coli* strains that were isolated concomitantly with *Enterococcus faecalis* after an open foot fracture caused by the 2004 tsunami resulting in a 10-year chronic bone and joint infection. Initially present antimicrobial resistant *E. coli* ST405 and ST940 isolates were followed by host adapted isolates of ubiquitous ST131 clone presumably acquired from the environment already upon initial foot fracture. The *E. coli* ST131 clade C1 strains showed genomic alterations associated with virulence and persistence including large chromosomal inversions and, subsequently, a large deletion to cause small colony variants and higher susceptibility to formaldehyde and other stress provoking In this context deletion of *hemB* catalyzing an early step in the pathway for heme biosynthesis was the major, but presumably not the only cause of small colony variant emergence. Surprisingly, ST131 isolates did not display pronounced biofilm formation in conventional biofilm assays suggesting unconventional modes of persistence. In summary, the genomes of ST131 clone members are highly plastic which enables their persistence in novel ecological niches. In individuals with underlying metabolic diseases such as diabetes wound infection can prepare for colonization with ST131 *E. coli* isolates.

**Funding:** This work was partially funded by ALF, the Petrus and Augusta Hedlunds Foundation and the Karolinska Institutet.

## INTRODUCTION

Being a ubiquitous commensal of the gastrointestinal tract of almost every human, isolates of the genetically diverse *Escherichia coli* can readily develop into a pathogen by few subsequently occurring genetic alterations, depending on the genomic context, by habitat switch or by overgrowth. For example, Enterotoxigenic *E. coli* (ETEC) arise by the acquisition of virulence plasmids and few genomic alterations ^1^. Also, the probiotic strain *E. coli* Nissle 1917 is closely related to the uropathogenic isolate CT087 ^2^, but the consequence of virulence factor expression is dependent on the genetic context ^3^ equally as virulence can be promoted by a habitat switch ^4^. Stress conditions and impaired immune responses can lead to microbial overgrowth and translocation ^5^

The extended use of antimicrobial agents has triggered the development of distinct multidrug resistant pandemic pathogenic clones. The *E. coli* sequence type ST131 is today the most widely distributed clone among fluoroquinolone and/or extended spectrum beta-lactamase (ESBL) resistant isolates world-wide ^6^.^7-9^. The development of ST131 clone members from antibiotic susceptible commensal isolates to pandemic highly prevalent multiresistant pathogens can be followed starting from the 19^th^ century with distinct genetic alterations and has accelerated in the 1980s/1990s. Thereby, three genetically clearly distinguishable clades A, B and C with distinct antimicrobial resistance patterns, alleles (single nucleotide polymorphisms), plasmids and accessory genomes can be discriminated, with a recent expansion of the multidrug resistant clades C1 and C2 ^6^.

The high-risk clone *E. coli* ST131 is predominantly associated with community as well as hospital acquired (nosocomial) urinary tract and blood stream infections in humans ^7^. Its success as a pathogen is also based on the fact that ST131 can also efficiently colonize the gastrointestinal tract with successful spread among and by asymptomatic community carriers and can be readily transmitted from person to person. Moreover, ST131 members can even cause disease in animals including wild and companion animals. Transmission can also occur by ST131 strains present in poultry, vegetable and other food. In the environment ST131 members can persist in wastewaters.

Although not exceptionally antibiotic resistant as compared to other *E. coli* clones such as ST405, the success of ST131 is partly based on its ability to sequentially acquire point mutations and genes that contribute to enhanced antimicrobial resistance. Early developed chromosomal resistance against fluoroquinolones has been accompanied by subsequently acquired ESBL genes CTX-M-14 (clade C1) and CTX-M-15 (clade C2) on different plasmid backgrounds and even on the chromosome.

Pathogens can adopt two fundamentally different virulence strategies leading either to an acute or a chronic infection. Expression of virulence factors leads to highly immunogenic acute infections readily tackled by antimicrobial therapy. The chronic infection status is associated with substantial metabolic adaptation which includes formation of biofilms, persister cells and small colony variants (SCV) ^10-12^. These metabolic variants grow slow on nutrient-rich medium under aerobic conditions compared to their otherwise fast *in vitro* growing ancestors or counterparts which cause acute infection. Already at the beginning of the 20^th^ century small colony variants of *Staphylococcus aureus* and subsequently *E. coli* had been recognized ^13^. In particular, *Pseudomonas aeruginosa* develops SCVs during chronic lung infection in patients with cystic fibrosis. The molecular basis for SCV development is auxotrophy for various classes of metabolites such as hemin, menachinone, thymidine, CO_2_ and lipoic acid with the most common physiological denominator being a defect in the electron transport chain and decrease in the proton motif force (PMF) that ultimately leads to increased resistance to aminoglycosides, but also susceptibility to antibiotics that are substrates of PMF-dependent efflux pumps ^14-19^. Consequently, exposure to aminoglycosides and other antibiotics, but also the immune system, antimicrobial peptides, the intracellular host environment and biofilm formation have been recognized as selective pressures that lead to selection for small colony variants *in vitro* ^20-24^. Nevertheless, small colony variants can have an advantage *in vivo* including being high biofilm formers ^11, 24, 25^. Small colony variants of *E. coli* have rarely been described in detail, although the phenotype has been reported to arise during gastroenteritis, urinary tract infection, bacteremia and chronic hip infection ^15, 26, 27^

Not mutually exclusive, adaptation mechanisms during chronic infections include the emergence of large chromosomal inversions, arisal of mutator strains and expansion of IS elements ^28^. Large chromosomal inversions are associated with speciation, their occurrence upon chronic infection indicates niche adaptation ^29-31^. Mutator strains, which lack repair of replication errors, facilitate adaptation, but might on the other hand lead to detrimental accumulation of mutations ^32-34^.

In this work, we describe the features of seven *E. coli* isolates that were recovered from a more than 10 year long wound infection with chronicity supported by the presence of foreign body material and sequester. With the multiresistant isolates to harbour IncFII based multireplicon plasmids with different versions of ESBL genes, the three persistent isolates were host adapted members of the pandemic clone ST131 with substantial chromosomal alterations such as large genome rearrangements and deletions which had developed a small colony variant phenotype.

## RESULTS

### Case description

An individual with a foot fracture caused during the 2004 tsunami suffered post-traumatically from a ten year long chronic bacterial tissue infection (Figure 1, Ahl et al, manuscript in preparation). While initially a diverse microbiological flora was observed, only multiresistant *E. coli* and an *Enterococcus faecalis* isolate persisted. We estimated *E. coli* to be the causative agent of the recurrent infection that was eventually cured in 2015 (Figure 1, Ahl et al, manuscript in preparation).

**FIG 1.**
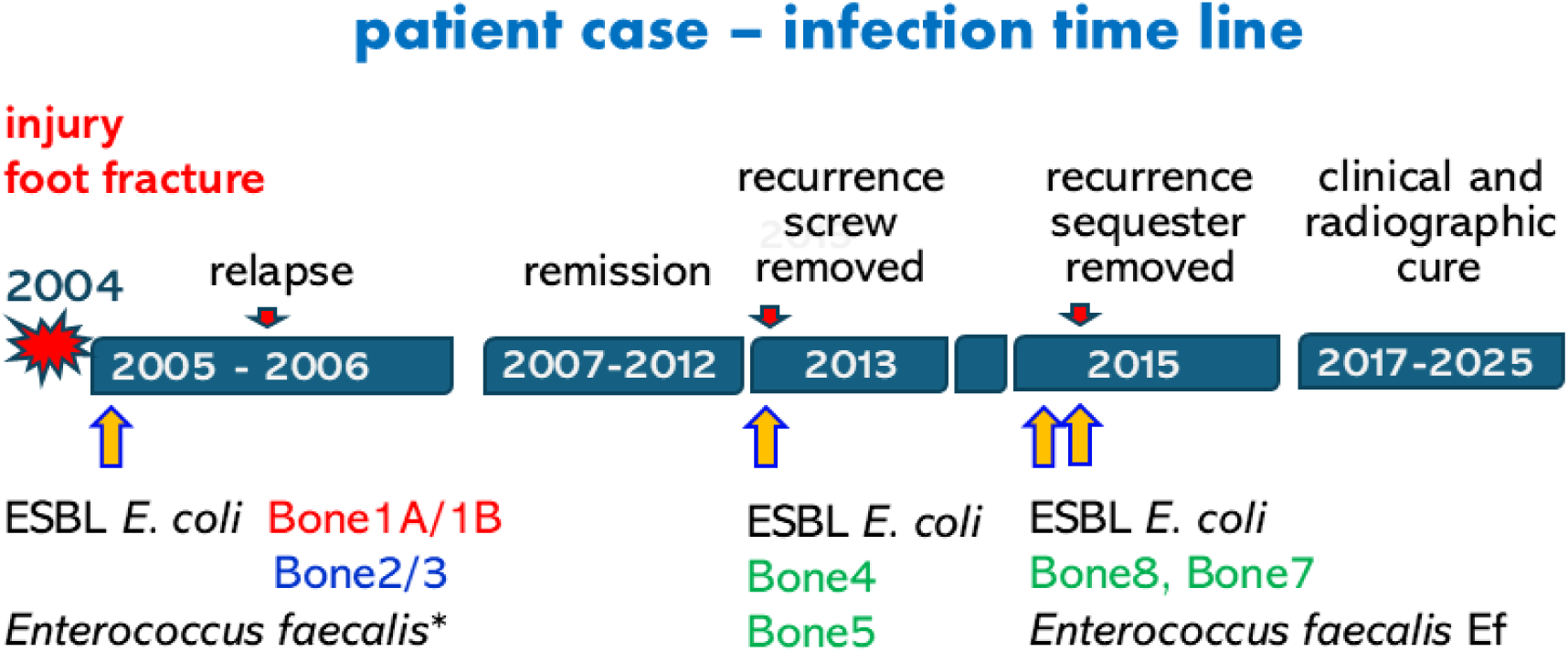
Timeline of the chronic infection after foot fracture caused during the 2004 tsunami. Red sign, injury December 2004; red arrow, relapse and measures such as removal of screws and sequesters. Orange arrow, recovery of bacterial isolates. Designation of *E. coli* strains is indicated; red=ST904; blue=ST405 and green=ST131. *, not maintained.

Although not being rare, persistent bone and tissue infections caused by isolates of the species *E. coli* are infrequently described^35, 36^. To unravel the molecular basis of adaptation and persistence, we analyzed the three *E. coli* isolates recovered at the beginning of the infection in 2005 (Bone1A and 1B (same isolation date with different colony morphologies on blood agar plates) and Bone2), two isolates from 2013 upon recurrence of the infection after more than seven year of silence (Bone4 and Bone5) and two more isolates - Bone8 and Bone7 – recovered in 2015 after two years of silence isolated within a three months interval in 2015 shortly before the infection was eventually cured (Figure 1; Supplementary Table 1).

### Phylogenetic placement of the isolates

Pulsed-field gel electrophoresis^37, 38^, and, after whole genome sequencing (see Material and Methods and Supplementary Material), *in silico* multilocus sequence typing (MLST) of seven genes^39^ and core genome phylogenetic assessment of the seven *E. coli* isolates initially classified the early isolates Bone1A,B and Bone2 (Bone3 displayed identical in macrorestriction fingerprinting and was not further analyzed) to possess distinct genetic backgrounds and, consequently, sequence types (Supplementary Table 2). In particular, Bone1A,B and Bone2 were sequence type (ST) ST940 and ST405, respectively, the latter known to be especially antimicrobial resistant^40^. The subsequent four isolates, Bone4,5,8,7 belonged to ST131, a pandemic *E. coli* clone predominantly causing urinary tract infection and bacteremia positioned in phylogroup B2. To further define the phylogenetic position of the Bone isolates, we selected representative *E. coli* ST131 isolates, and used them to define *E. coli* phylogenies based on core-genome alignment. While ST131 isolates formed a congruent cluster, Bone1A,B and Bone2 are positioned on a long branch (Supplementary Figure 1). Subsequently, compared to representative strains from the different *E. coli* phylogroups ^41^, Bone1A,B were phylogroup B1 and Bone2 phylogroup D genetically distinct from later collected ST131 isolates Bone4,5,8,7 (Figure 2A).

**FIG 2.**
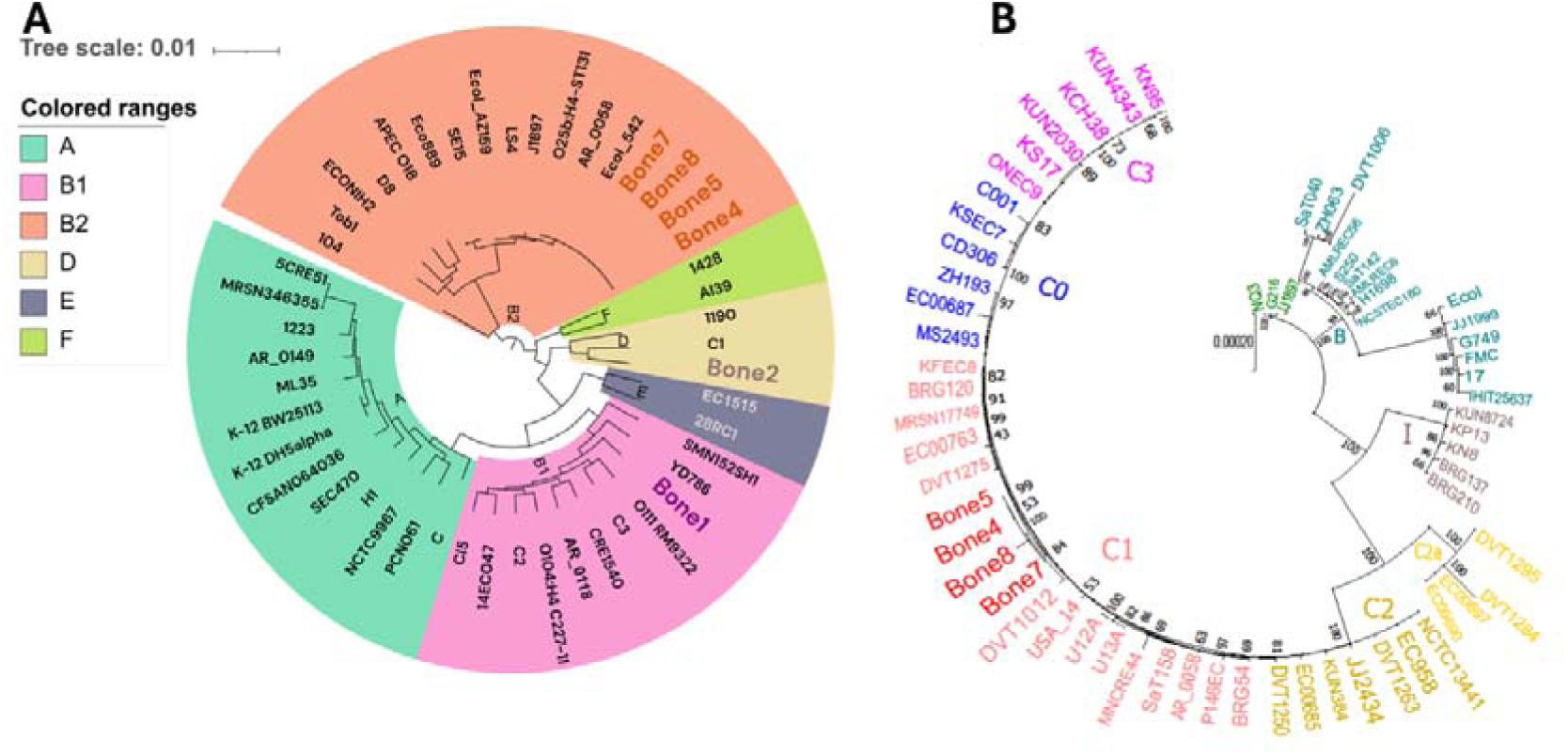
Phylogenetic classification of the *E. coli* Bone isolates. (A The maximum likelihood phylogenetic tree based on the core genome alignment of 42 strains selected to represent the six established *E. coli* phylogroups A, B1, B2, D, E and F classified Bone1 isolates in phylogroup B1, the Bone2 isolate in phylogroup C and Bone4,5,8 and 7 isolates in phylogroup B2. B) Phylogenetic classification of the ST131 Bone isolates within ST131 isolates. The Maximum likelihood phylogenetic tree based on the core genome alignment of ST131 *E. coli* clades B, C0, C1, C2 isolates showing the positions of Bone4,5,8 and 7 isolates within the ST131 strains of clade C1.

Pandemic ST131 pathogens have diversified into three major clades with a recent expansion of clade C into multiresistant subclades C1 and C2. To further position the Bone isolates within the ST131 clonal population, ST131 strains of different major clades were chosen to construct a core-genome based phylogenetic tree. Bone4/5/8/7 constitute a dense group of closely related strains within clade C1 strains (Figure 2B, Supplementary Figure 1). The basic genomic characteristics of the strains are summarized in Supplementary Table 2.

### Altered chromosomal targets and acquired genes confer the antibiotic resistance profile

All strains were multidrug resistant with experimentally determined resistance against the fluoroquinolone ciprofloxacin and the third-generation class cephalosporin cefotaxime (Supplementary Figure 2A). Bone1A,B and 2 were also resistant against the third-generation class cephalosporin ceftazidime. Furthermore, all strains except Bone2 displayed resistance against the aminoglycoside gentamicin, and, except Bone8 and Bone7, were resistant against trimetroprim/sulfamethoxazole.

Experimentally determined resistance was grossly consistent with mutations in relevant targets and acquired resistance genes (Supplementary Figure 2B, C and 3). Notable were the amino acid substitutions characteristic for mediating fluoroquinolone resistance present in the gyrase subunit GyrA, S83L D87N and in the DNA topoisomerase IV subunit ParC S80I in all isolates.

Distinct, but related, large multireplicon IncFII plasmids harbor the majority of acquired antimicrobial resistance determinants in all strains (Supplementary Table 2; Supplementary Figure 2 and 3A and C)^42, 43^. While the four origin of replication IncFII type F36:F22:A1:B1 plasmid of Bone1A,B harbors a broad-spectrum CTX-M-15 beta-lactamase (consistent with activity against third-generation cephalosporines cefotaxime and ceftazidime)^44^, a broad-spectrum CTX-M-14 beta-lactamase characteristic for ST131 clade C1 strains are present on the four origin of replication IncFII plasmids of type F1:A2:B20/Col156 of Bone4-8 (Supplementary Table 2: Supplementary Figure 2C). Of note, the IncFII plasmids of the ST131 strains Bone8 and Bone7 have otherwise lost the majority of antimicrobial resistance determinants (Supplementary Figure 2C and 3A). All plasmids are composite plasmids with the Bone IncFII plasmids to possess long highly similar regions such as the conjugative transfer system and have highly similar homologs in the database (Supplementary Figure 3C and D).

**FIG 3.**
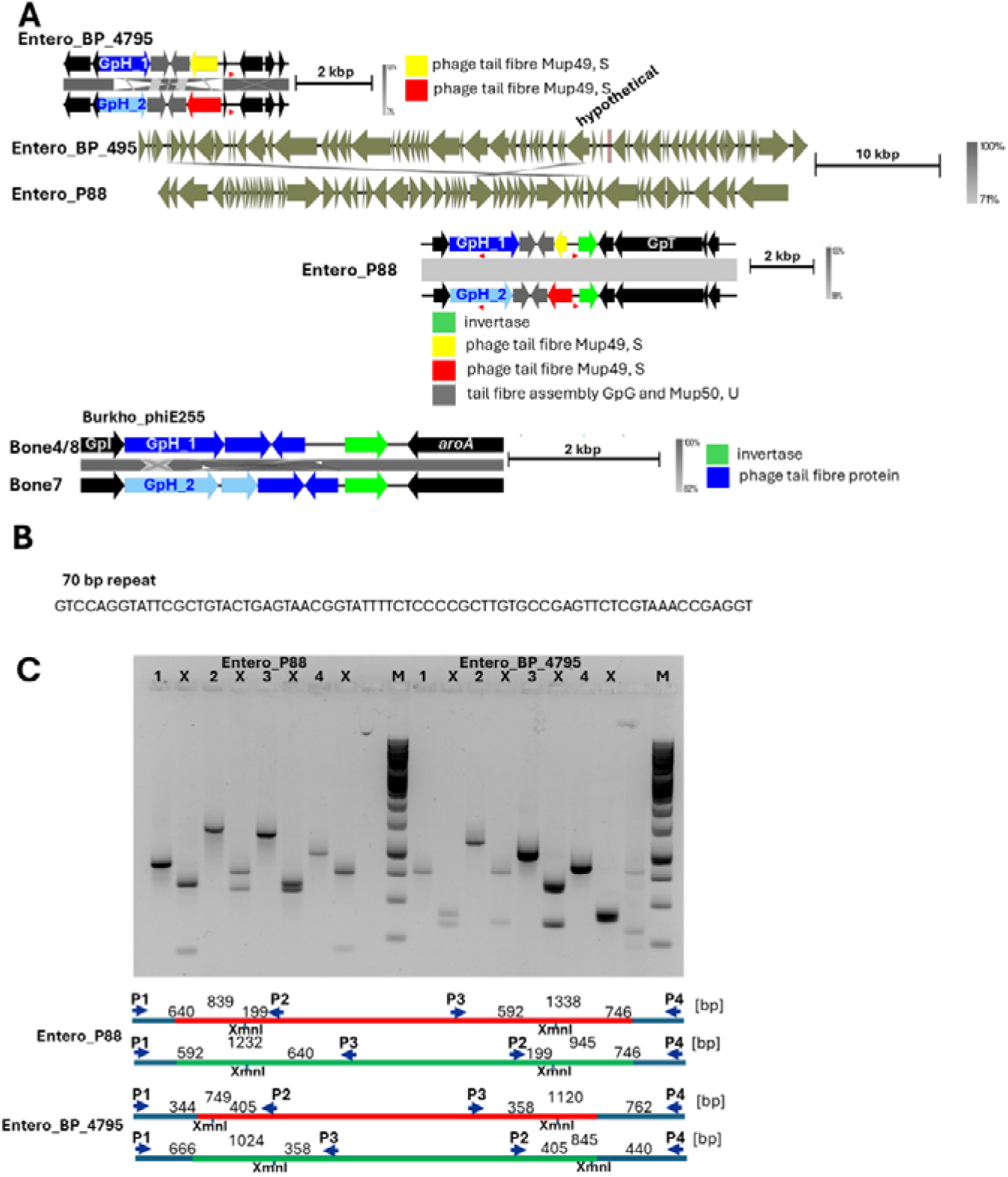
Entero_BP_4795, Entero_P88 and Burkho_phiE255 phages displaying DNA inversions or translocation to alter the sequences of homologous tail fiber GpH and tail fiber Mup49, S proteins. A Genomes of Entero_BP_4795, Entero_P88 and Burkho_phiE255 phages showing the homologous regions, observed DNA inversions and resulting protein variants. The impact of the DNA inversions is unclear. Red arrows, 70 bp direct repeats. B Nucleotide sequence of the 70 bp direct repeat. C PCR analysis and XmnI digest to confirm the switches in the DNA fragments in Entero_P88 and Entero_BP_4795 phages using Bone4 as a template. PCR fragments and restriction digests were separated by 1% agarose gel electrophoresis in 1x TAE buffer. 1-4 indicates PCR fragments in respective order in the inversion region. X=XmnI digest.

Additional resistance determinants are encoded on other plasmids in Bone1 and Bone2, on the chromosome including a copy of the CTX-M-15 beta-lactamase in Bone2 and a second copy of the CTX-M-14 beta-lactamase in Bone4-8 (Supplementary Table 2; Supplementary Figure 2B, C and 3B). In summary, a broad panel of acquired resistance genes and chromosomal mutations actually and potentially mediating antimicrobial resistance has been originally present in the *E. coli* Bone strains.

### Initial characterization of ST131 Bone isolate genomes

The ST131 representatives had first been recovered eight years after onset of the chronic wound infection. We investigated whether the chromosomes of ST131 Bone isolates harbored specific virulence factors or other characteristics which could indicate mechanisms of success in chronic tissue infections compared to the early *E. coli* Bone1A,B and Bone2 isolates and a reference ST131 clade C1 isolate. Before starting the analysis, we normalized the chromosomes to the origin of replication (ORI) which is located between the *mioC* and *gidA* genes in *E. coli* taking the first nucleotide of the first high affinity DnaA binding site in clock-wise direction as a reference (Supplementary Figure 4; ^45, 46^). All genomes consisted of one circular chromosome, one size-variable multiORI IncFII plasmid and a variable number of additional plasmids (Supplementary Table 2, Supplementary Figure 3A, B and D). Chromosome size varied between 4.823 Mbp (Bone1) and 5.167 Mbp (Bone4).

All Bone strains lack the type III secretion system as a major virulence factor, but code for a type VI secretion system, a common microbial, and, occasionally eucaryotic, warfare system ^47, 48^, the inverse autotransporter intimin-like adhesin FdeC (and the two glutamate decarboxylases as core genome virulence genes mediating stress resistance) (Supplementary Figure 5). The IncF plasmids harbor the SenB enterotoxin and a microcin M bacteriocin.

Virulence factors specific for ST131 strains include the Entero_lambda phage encoded *iss* for increased serum survival, three type 5 secretion autotransporter SPATE (Serine Protease Autotransporters of Enterobacteriaceae) proteases and the SinHI RatA invasins (also present in Bone2). The ST131 isolates harbor the KpsMII-K5 capsule type reported to be associated with enhanced virulence ^49^, while Bone2 contains the KpsMIII-K98 capsule type (complete list of identified virulence factors in Supplementary Figure 5A). The ST131 isolates and Bone2 code for three siderophore biosynthesis and transport gene clusters, enterobactin, aerobactin and yersiniabactin, while ST131 isolates code for the largest number of iron uptake systems (Supplementary Figure 5B and C).

Nine complete prophages (predicted) are integrated into the ST131 *E. coli* chromosomes, while Bone1A,B and Bone2 harbor one and five complete prophages, respectively (Supplementary Figure 5D). Of note is phage mEp460 which occurs in two distinct copies in the Bone4-8 genomes (Supplementary Figure 5E). Identified complete prophages belonged to the class of Caudoviricetes, family Peduoviridae and the genera Peduovirus, Lambdavirus, Bcepmuvirus, Marienburgvirus, Xuanwuvirus and Gegevirus (Bone2). In addition, phages whereby functionality cannot be predicted with certainty and incomplete phage(s) were identified integrated into all Bone isolate chromosomes (Supplementary Figure 5D).

Of note, prophage sequences were subject to directed reversible DNA recombination (switch) within one isolate (Figure 3 and Supplementary Figure 5F). Reversible inversion events of DNA segments were identified in Entero_P88 and Entero_BP-4795 between the phage tail fibre protein GpH and the phage tail fibre protein Mup49, S in all ST131 Bone isolates leading to alterations in the amino acid sequence of the two proteins (Supplementary Figure 5G). Further more, a DNA translocation has been identified in Burkho_phiE255 in Bone7 compared to Bone4 and Bone8 isolates again leading to alterations in the amino acid sequence of the phage tail fibre protein GpH and the phage tail fibre protein Mup49, S that are homologous between the three phages Enterp_P88, Entero_BP-4795 and Burkho_phiE255. Of note, two of these prophages, Entero_P88 and Burkho_phiE255 contain a phage DNA invertase (Figure 3A). Direct repeats of 70 bp flanking the inverted region have been identified in Entero_P88. The same repeat sequence occurs also once in Burkho_phiE255 at the translocation site (Figure 3). Furthermore, prophage insertion can alter the host physiology and regulation in multiple ways such as harboring virulence factors (Supplementary Figure 5A). We have observed that some of the phages are not inserted downstream of tRNAs (example in Supplementary Figure 5E).

Flagella-mediated swimming and surface-induced swarming motility are integral parts of the acute as well as chronic infection process equally as biofilm formation ^50, 51^. As only few *E. coli* clones, ST131 including the Bone isolates possess two types of flagellar regulons, the conventional peritrichous Flag-1 system and an ancient lateral Flag-2 system at alternative chromosomal locations (Supplementary Figure 6A). In vitro, only the Flag-1 system contributes to flagella mediated swimming and swarming motility, while the role of the lateral Flag-2 system is unknown ^52^.

As chronic infections are conventionally associated with (enhanced) biofilm formation, we assessed whether extracellular appendages commonly involved in biofilm formation such as proteinaceous type 1 fimbriae, amyloid curli, the *E. coli* common pilus and Antigen 43 (Ag43) and exopolysaccharides such as cellulose, poly-N-acetylglucosamine (PNAG) and colanic acid are present in the Bone isolates (Supplementary Figure 6A-E). Of note, type 1 fimbriae display the ST131 allele variant which has an integrated ISEc53 element disrupting *fimB* encoding a recombinase involved in phase-dependent expression, while Bone 1A,B is lacking type 1 fimbriae. ST131 Bone4 and 5 possess two Ag43 loci, while one of those is inactivated in Bone8 and 7. Bone two possesses four Ag43 loci with the expansion of Ag43_1. The PNAG operon, which can be variable in presence and location in *E. coli* ^2^, is present at the same location in all strains as in all ST131 isolates and the *E. coli* K-12 MG1655 reference. The Bone2 isolate has a large insertion close to the PNAG operon.

Cyclic di-GMP is a ubiquitous second messenger regulating the lifestyle switch between sessility and motility in Bacteria ^53^. The cyclic di-GMP signaling system is highly plastic and variable thus flexibly regulating biofilm formation versus motility ^54, 55^. The majority of cyclic di-GMP turnover proteins, GGDEF diguanylate cyclase and EAL phosphodiesterase and their evolved domains which lack catalytic activity harbor specific, often multiple, amino acid substitutions and occasionally N- or C-terminal extensions/deletions compared to the reference ST10 *E. coli* K-12 MG1655/Fec10 ^56^ (Supplementary Figure 7A; Supplementary Table 3). In this context, cyclic di-GMP turnover proteins of the ST131 Bone isolates are highly similar to their ST131 representatives from the different clades. Already individual amino acid substitutions, however, can significantly alter the in vivo activity of these proteins ^2, 55, 57^. Furthermore, as major cyclic di-GMP network alterations, ST131 Bone isolates, as all ST131 reference isolates, harbor a YcgG protein truncated for the N-terminal CSS signaling domain and had transformed the YddV/DosC GGDEF domain protein into a novel short protein (Supplementary Table 3, Supplementary Figure 7B and C)

### Genome rearrangements between Bone4 and the representative clade C1 isolate AR_0053

We were subsequently wondering whether ST131 Bone isolates, as exemplified with Bone4, contained specific genome characteristics compared to a representative clade C1 isolate. AR_0058 was chosen for comparison as this isolate has a completely sequenced genome (CP021689.1). Eight regions of difference >20 kbp were found, whereby six regions represented insertions in Bone4 and one exchange of DNA sequences between Bone4 and AR_0058 (Supplementary Table 4A). Phages ΦAA91-ss, mEP460 (2x), Shigell_SfII and TL-2011b are present in Bone4, but not AR_0058 (Supplementary Figure 5D and Table 4A and Ffi). Furthermore, a 67211 bp long chromosomal region with genes coding for the general type II secretion system, Ag43 and the polysialic acid capsule K1 and a 21428 bp long region is present in Bone4, but absent in AR_0058. This chromosomal region is subject to variability in ST131 isolates as genes encoding Ag43, the K1 capsule, but also the aerobactin siderophore is absent in clade A strain SE15 (data not shown). Present in AR_0058 and absent in Bone4 is the Lederbergvirus Shigel_Sf6/Entero_Sf101.

Among the <20 kbp regions of difference were the transposition of the class A beta-lactamase CTX-M-14 colocalized with different mobile element genes leading to a second chromosomal copy of this resistance gene in Bone4 and the other ST131 Bone isolates (Supplementary Figure 3C). This transposition disrupted the y2843 protease. Furthermore, transposition of IS1 family members has occurred located predominantly in intergenic regions.

In addition, single nucleotide deletions and polymorphisms (SNPs) were observed. Multiple single nucleotide deletions led to the restoration of pseudogenes in Bone4 (Supplementary Table 4B). The consequences of synonymous and non-synonymous SNPs, several of them unique to ST131 Bone isolates need to be further investigated.

### Major genome rearrangements within ST131 Bone strains

#### ST131 Bone isolates harbour large chromosomal inversions

The chromosomes of representative strains of the major ST131 clades A, B, C1 and C2 are in synteny with the chromosome of the ST10 reference strain *E. coli* K-12 MG1655. Comparison of the chromosomes of Bone4-8 with the ST131 reference genomes showed distinct large scale genome rearrangements caused by homologous recombination between two rRNA operons for Bone4, 5 and 7 (Figure 4A). Considering the first high affinity binding site for DnaA at the origin of replication as reference (Supplementary Figure 4), the >3.5 Mbp inversion of the Bone4 chromosome caused a substantial replichore dissymmetry, while the inversion of almost the entire chromosome observed in Bone7 does not strongly impair the position of the terminus of replication.

**FIG 4.**
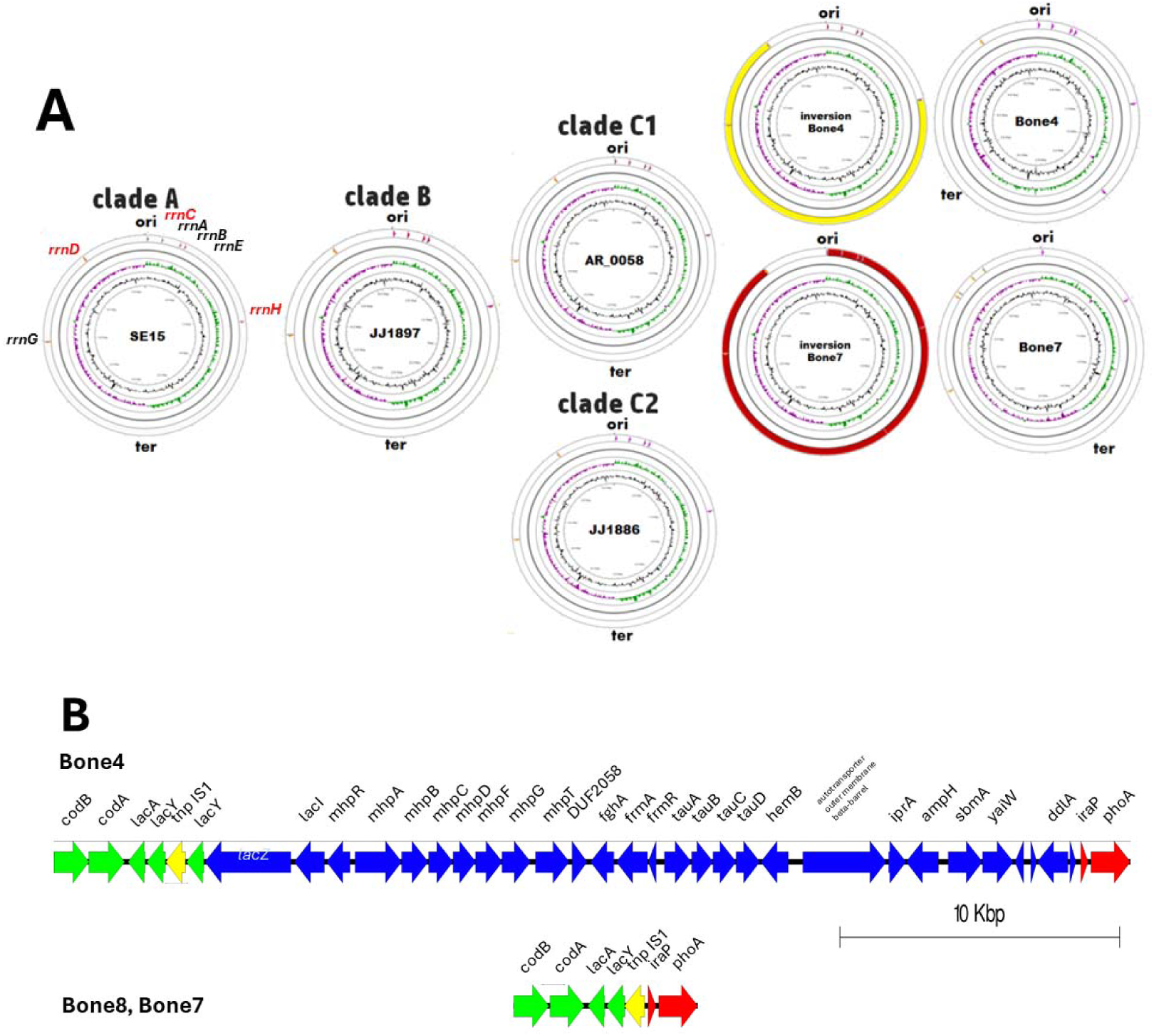
Major chromosome alterations in and among ST131 isolates. A The chromosomes of Bone4. Bone5 and Bone7 isolates possess large chromosomal inversions compared with the chromosomes of representative isolates of the major ST131 clades. In Bone4 (Bone5 equal to Bone4, not displayed), the recombination between the rRNA operons leads to a large unbalanced inversion around *oriC* yielding an asymmetric chromosome with the terminus of replication (*terR*) shifted as indicated by the G+C skew of the leading strand. In Bone7 almost the entire chromosome is inverted taking the origin of replication (ori) as the reference. While the chromosome remains symmetric, the recombination between two rRNA operons led to a change in the gene position between the leading and lagging strand for almost all the genes. The G+C skew and ribosomal RNA operons are indicated by red arrows in the outermost ring. All reference strains of clades A, B, and C are colinear and seven rRNA operons arranged around the origin of replication comparable to *E. coli* K-12 MG1655. Five rRNA are transcribed clockwise and two rRNA operons are transcribed counter-clockwise. the G+C content are shown in the inner circles. Reference strains for the ST131 clades *E. coli* SE15, clade A (not displayed); *E. coli* JJ1897, clade B; *E. coli* AR_0058 and MRSN17749, clade C1 and *E. coli* JJ1886, clade C2. B A major deletion of 32922 bps comprising the core genome region between *lacY* and *iraP* is present in ST131 isolates Bone8 and Bone7 compared to Bone4 and Bone5.

#### A large chromosomal deletion occurs in late Bone ST131 isolates

Besides large chromosomal inversions, additional genome alterations were observed. Indeed, Bone8 and Bone7 harbor one deletion of 31.9 kbp compared to Bone4 (Figure 4B). This deletion spans the core genome genes between *lacY* and *iraP* and includes the entire operon for the 3-(3-hydroxyphenyl)propionate degradation pathway *mhpRABCDFT*, the entire *tauA-D* taurine uptake and catabolism operon and *hemB* encoding the delta-aminolevulinic acid dehydratase catalyzing the early reaction in the tetrapyrrole biosynthesis eventually resulting in heme and siroheme biosynthesis. Other deleted genes include reported stress resistance genes and *ampH* involved in peptidoglycan remodelling and recycling.

Besides the 31.9 kbp deletion, distinct genomic alterations occur between Bone4 and Bone5, Bone7 and Bone8 strains. These include IS1 element transpositions, deletions/insertions and SNPs (Supplementary Table 4C and D). Of note is a 286 bp deletion in Bone7 which resulted in a novel fatty acid hydroxylase family fusion protein created from the open reading frames of the linoleoyl-CoA desaturase and a fatty acid hydroxylase family protein.

### Phenotypes and molecular basis of development of the sequential Bone isolates

#### Bone 8 and 7 strains display a small colony variant phenotype recovered by red blood cell lysate

We subsequently investigated whether the Bone isolates display differences in behavior, metabolism and physiology that would be reflected by the genome analyses. Growth on different agar media showed that Bone1A,B and Bone2 and the ST131 isolate Bone4 grew well on all media. ST131 isolates Bone8 and Bone7 grew similar as Bone4 on blood and Mueller Hinton Fastidious agar, but a small colony variant colony phenotype was observed on Mueller Hinton and LB agar consistent with the deletion of *hemB* (Supplementary Figure 8 A and data not shown). All ST131 strains showed the common lactose negative phenotype for ST131 isolates on CLED agar (Supplementary Figure 8B). Partial growth restoration occurred upon provision of hemin (factor X), but not nicotinamide-adenine-dinucleotide (NAD; factor V) on Mueller Hinton agar plates; auxotrophies for both factors are commonly associated with host adaptation of microbes (Supplementary Figure 8C).

We assessed the growth of the strains quantitatively in liquid medium under low oxygen pressure in the 96-well plate assay (Figure 5 and 6A). The logarithmic growth was almost equal for all isolates. Bone8 and Bone7 isolates, however, showed lower optical density in the stationary phase of growth in Mueller Hinton, Bolton and LB broth with relative OD most similar in Bolton broth and most different in LB medium. Addition of hemin led to a growth recovery of Bone7. Of note, supplementation with 5% lysed red blood cells led not only to recovery of steady state biomass in Bone8 and Bone7 isolates, but triggered significantly enhanced growth and higher optical density in the stationary phase of growth for all investigated isolates displaying almost identical growth curves (Figure 5D and E). This indicates that required metabolites and carbon sources for optimal growth are provided by the red blood cell lysate.

**FIG 5.**
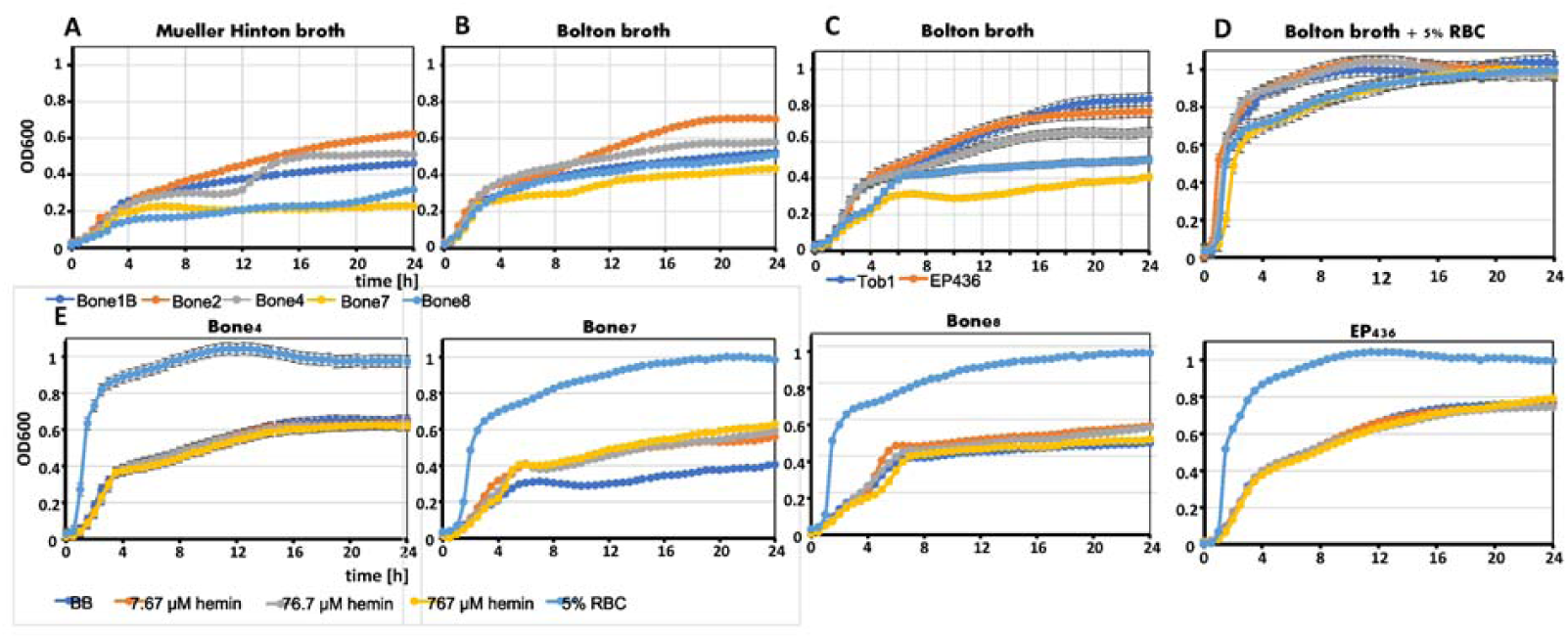
Growth characteristics of Bone isolates and reference strains in liquid medium. A Comparative assessment of growth of Bone isolates in Mueller Hinton broth. B Comparative assessment of growth of Bone isolates in Bolton broth. C Comparative assessment of growth of Bone isolates and reference strains (commensal E. coli TOB1 (phylogroup B2) and ST131 EP436 strain (blood isolate) in Bolton broth. D Comparative assessment of growth of Bone isolates and reference strains in Bolton broth with 5% lysed red blood cells (RBC). E Growth of Bone isolates and reference ST131 EP436 strain in Bolton broth, Bolton broth with 7.67, 76.7 and 767 µM hemin and Bolton broth with 5% lysed RBC. Ten µl of a cell suspension of OD=1 was inoculated in 200 µl culture medium in individual wells of a 96-well plate and the plate was incubated at 37° C and OD_600_ measured in a *SpectraMax i3x* (Molecular Systems). Displayed is one of two representative experiments performed in duplicate.

**FIG 6.**
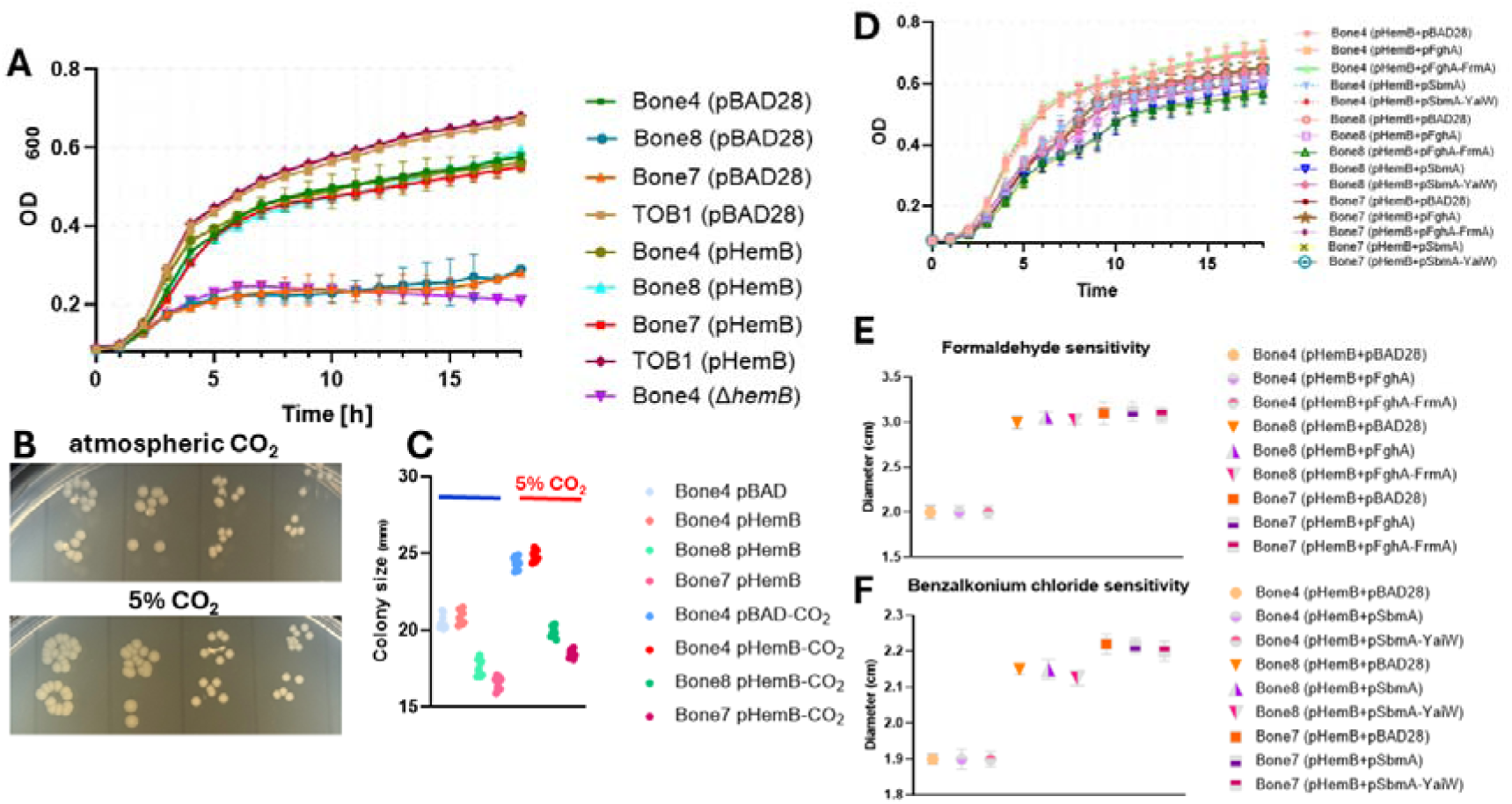
Complementation of small colony variant and susceptibility phenotypes of Bone8 and Bone7 isolates and creation of a *hemB* deletion in Bone4. A Growth characteristics of Bone4, 8 and 7 pBAD28 vector control compared to Bone4, 8 and 7 *hemB* complemented strains and Bone4 Δ*hemB* in LB medium. Commensal *E. coli* TOB1 (phylogroup B2) served as a reference isolate. B Colonies of Bone4 pBAD28 vector control compared to Bone4, 8 and 7 *hemB* complemented strains grown on LB agar plates under atmospheric and 5% CO_2_ pressure at 37 °C. C Quantitfication of colony diameter for growth on LB agar plates as shown in B. D Comparative assessment of growth for Bone4, 8 and 7 isolates after combined complementation with *hemB* on plasmid pSRKTc and FghA (FrmB), FghA-FrmA, SmbA and SmbA-YaiW. FghA-FrmA and SmbA-YaiW located in the 31922 bp region deleted in Bone8 and Bone7 have been reported to be involved in resistance against formaldehyde and quarternary ammonium compounds, respectively ^58, 59^. Strains were grown in LB medium at 37 °C. E Complementation of pHemBTc bearing Bone4, 8 and 7 isolates with pFghA and FghA-FrmA to assess the effect on formaldehyde sensitivity by the disk assay. Strains were grown on an LB agar plate at 37 °Cwith the inhibition zone assessed after 24 h of growth. pHemBTc=*hemB* cloned in pSRKTc; pFghA and pFghA-FrmA=*fghA* and *fghA-frmA* cloned in pBAD28 (Supplementary Table 6). F Complementation of pHemBTc bearing Bone4, 8 and 7 isolates with pSmbA and pSmbA-YaiW to assess the effect on formaldehyde sensitivity by the disk assay. Strains were grown on an LB agar plate at 37 °Cwith the inhibition zone assessed after 24 h of growth. pHemBTc=*hemB* cloned in pSRKTc; pSmbA and pSmbA-YaiW=*smbA* and *smbA-yaiW* cloned in pBAD28 (Supplementary Table 6).

Subsequently, we tested whether the persistent ST131 Bone isolates developed resistance towards stress factors relevant in the tissue environment (Supplementary Figure 9). All isolates displayed equal sensitivity to bleomycin, thiocyanate, ascorbic acid and formic acid, but Bone8/7 isolates were significantly more sensitive towards formaldehyde and gradually against benzalkonium chloride, while slightly more sensitive towards rifampicin and H_2_O_2_.

The >30 kbp deletion encompasses *hemB* that can be causative for the SCV phenotype. Impairment of heme biosynthesis impairs multiple metabolic functions such as the respiratory chain and the defense against oxygen radicals as heme is a co-factor in enzymes (Supplementary Figure 8D). We therefore cloned *hemB* into the pBAD28 expression vector and assessed its effect on colony size and growth. Expression of *hemB* (already without inducing the expression by L-arabinose) almost fully recovered the SCV phenotype of Bone8 and Bone7, while it had a minor growth enhancing effect on Bone4 and no effect on the commensal isolate TOB1 included as a phylogroup B2 reference strain (Figure 6A). Growth of the complemented strains on LB agar plates showed significant, but not full recovery of the colony size under atmospheric conditions and when grown under 5% CO_2_ (Figure 6B and C). Of note, colony size has been observed to be proportionally dependent on high CO_2_. While *hemB* is a major determinant of growth and colony size recovery, the experimental set-up will allow the identification of additional genes contributing to the SCV phenotype.

Other genes deleted include *sbmA-yaiW* reported to be involved in resistance against quaternary ammonium compounds including benzalkonium chloride ^58^ and the *frmR-frmA-fghA*(*frmB*) operon with *fghA* reported to contribute to formaldehyde detoxification ^59^ and *frmA* to oxidative stress resistance. As Bone8 and Bone7 strains were significantly more susceptible to the detergent benzalkonium chloride and formaldehyde, the contribution of *sbmA* and *fghA*, respectively, to intrinsic resistance was investigated (Figure 6D-F). In order to carry out the assay on standard agar plates and not on blood agar plates, all strains were investigated with *hemB* expressed from the pSRKGm vector. Of note, *hemB* complementation did not alter susceptibility (data not shown). However, expression of either *smbA*, *smbA-yaiW*, *fghA* and *frmA-fghA* altered the susceptibility against benzalkonium chloride and formaldehyde, respectively. We conclude that the enhanced susceptibility of Bone8 and Bone7 isolates might be multifactorial and/or entirely dependent on alternative gene products.

#### Biofilm formation ability of Bone isolates

Biofilm formation is known to be a major determinant for the chronic infection process (Costerton, Stewart et al. 1999; Römling and Balsalobre 2012). Among the biofilm phenotypes of *E. coli*, most ancient and conserved is the rdar morphotype composed of amyloid curli fimbriae and the exopolysaccharide cellulose and a poly-N-acetyl-glucosamine based biofilm ^2, 60, 61^. Assessment of the ability of the isolates to express rdar or PNAG-based biofilm indicated that the ST131 isolates Bone4,5,8,7 gradually showed enhanced dye binding which is delayed at 37°C and abolished upon addition of 11 mM glucose mimicking diabetic tissue (Figure 7A and B). Of note, dye binding can alternatively indicate increased outer membrane permeability in case of Bone8 and 7. Bone1A,B and Bone2 isolates did not display any dye binding. In Bone4, stimulation of biofilm formation by overexpression of the potent diguanylate cyclase AdrA demonstrated the capacity for high biofilm formation (Figure 7C).

**FIG 7.**
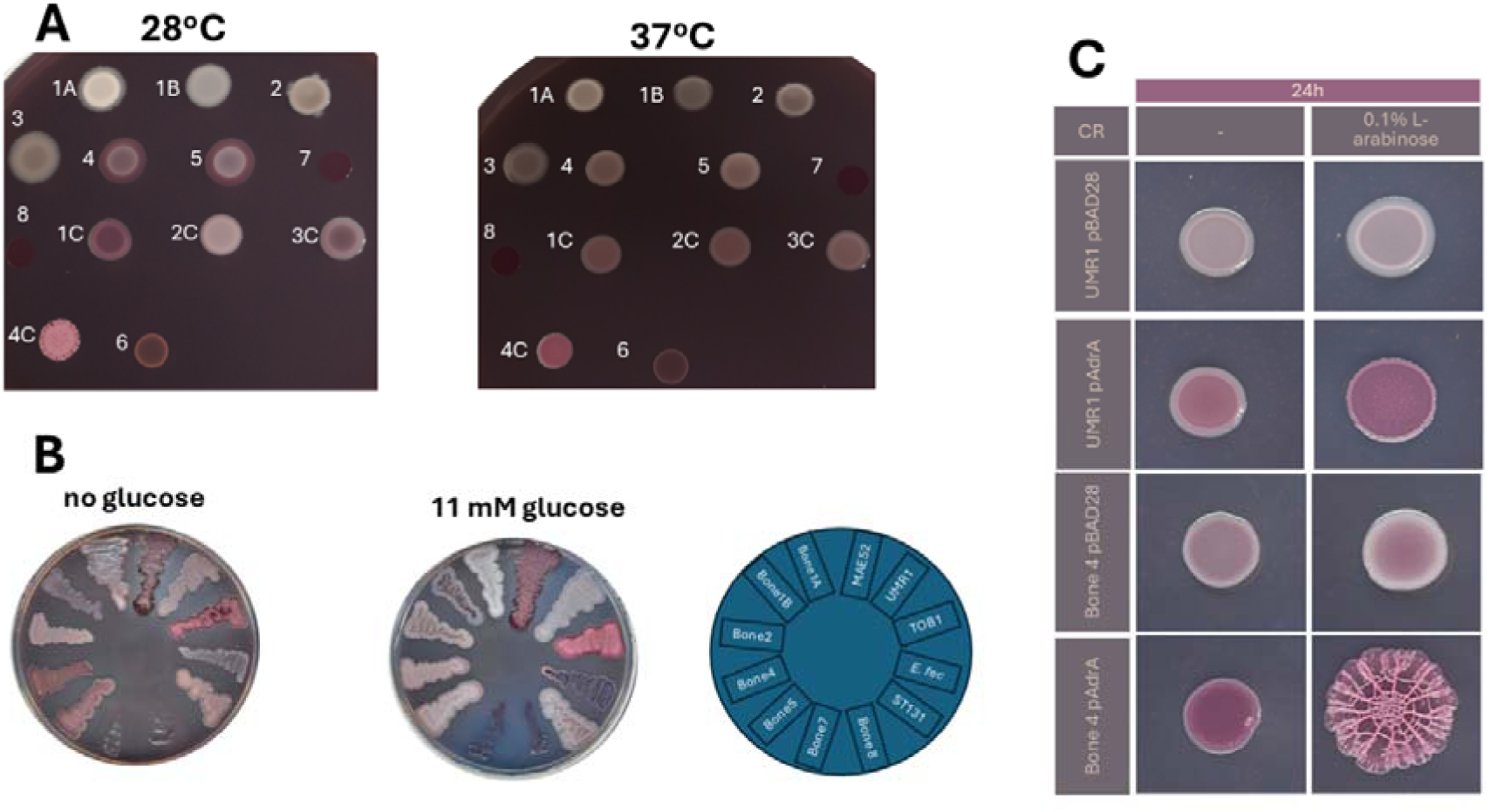
Rdar colony biofilm formation of Bone isolates. A Biofilm formation on Congo red LB without salt agar plates as displayed as the rdar morphotype. Designation of *E. coli* Bone isolates according to their numbers. Reference strains, 1C*, E. coli* TOB1 wild type control expressing curli fimbriae and cellulose; 2C, *E. coli* TOB2 (TOB1 Δ*csgD*), 3C, *E. coli* TOB2 (curli only expressing, TOB1 Δ*bcsA*), 4C, cellulose only expressing *E. coli* ECOR31 Δ*csgBA*. Growth for 72 h at 28 °C and 37 °C. B Rdar biofilm formation on Congo red LB without salt agar plates without and with 11 mM glucose. Growth for 72 h at 37 °C. C Rdar biofilm formation capacity of the *E. coli* ST131 isolate Bone4 compared to *S. typhimurium* UMR1 upon overexpression of the diguanylate cyclase AdrA induced by 0.1% L-arabinose. Growth for 24 h at 37 °C.

Alternatively, we assessed biofilm formation in liquid medium onto the abiotic polystyrene surface under conditions of type 1 fimbriae and rdar morphotype dependent biofilm formation (Supplementary Figure 10A). Biofilm formation was highest for *E. faecalis* with all the *E. coli* Bone isolates to display low biofilm formation under both growth conditions. Complementation with *hemB* did not significantly alter the biofilm forming ability. Coincubation with *E. faecalis* did not have an additive effect on biofilm formation (Supplementary Figure 10A). As *E. faecalis* has been recovered consistently from the wound, biofilm formation was also investigated in TSB + 1% glucose medium reported optimal for biofilm formation of *E. faecalis* isolates ^62^. Equally biofilm formation in relevant cell culture medium William’s Eagle and αMEM medium (with low (5 mM) and high (25 mM) glucose of all Bone isolates has been minimal (Supplementary Figure 10B). Surprisingly, all strains (besides the high biofilm positive control *S. typhimurium* MAE52), showed minimal biofilm formation that was not significantly enhanced upon coincubation with *E. faecalis*.

### Alterations in the regulons for peritrichous and lateral flagella

Assessment of motility in low density Bolton broth agar showed fast swimming motility of early isolates Bone1 and Bone2. However, also ST131 isolates Bone4, Bone8 and Bone7 showed residual swimming motility after subsequent extension of the assay time (Supplementary Figure 10C).

## DISCUSSION

*E. coli* ST131 and eventually *E. faecalis* have been causative agents of a chronic bone and joint infection that was eventually cured after a more than 10-year infection. While chronic tissue infections with *E. coli* are rarely investigated, in this work we have characterized the genomic features with a focus on the persistent *E. coli* ST131 strains as compared to initially isolated multidrug resistant *E. coli* strains that have not been maintained. Chronic infections are associated with genomic adaptations of microbes to ensure persistence in the host environment which can lead to substantial alterations in the genome ^30, 63^.

Indeed, the ST131 Bone4 strain isolated nine years after the onset of the chronic infection and seven years after the infection was initially silenced and subsequently Bone8 showed host adaptation by substantial genomic alterations in the form of a distinct large chromosomal inversion. Large chromosomal inversions have been shown to accompany chronic infections, host adaptation and speciation. For example, chromosomes of *P. aeruginosa* during chronic lung infection of cystic fibrosis patients and human host adaptation within *Salmonella enterica* from *Salmonella typhimurium* to *Salmonella typhi* is accompanied by large chromosomal inversions ^31, 63^. Those inversions occur between homologous sequences such as among rRNA operons and IS elements and can cause significantly asymmetric chromosomes with displacement of the terminus of replication (Figure 4A, ^64^). Thereby, large chromosomal inversion can modify the mode and direction of chromosomal evolution. It will certainly be of interest to assess the transcriptome and metabolome of those host-evolved Bone strains in order to get deeper insights into the altered cell physiology. However, the presumed heterogeneity and diversity of the isolates in the population causing the chronic wound infection could not been assessed as only one isolate per sampling point had been preserved.

A hallmark of the late ST131 Bone isolates has been their SCV phenotype. In clinical SCVs, one single mutation, though in different pathways, is causative for this phenotype. Indeed, in the context of a >30 kbp deletion, in the Bone8 and Bone7 strains, deletion of *hemB* is the major, but not the only cause contributing to the SCV phenotype. The complementation of the SCV phenotype with *hemB* will enable to detect additional genes involved in the SCV phenotype. The SCV phenotype was associated with enhanced formaldehyde and quarternary ammonium compound susceptibility of the isolates (Figure 6E and F; Supplementary Figure 9). However, FghA, which redundantly with YeiG detoxifies formaldehyde in *E. coli* ^59^, equally as the FrmA-FghA combination did not complement the formaldehyde susceptiblity. Additional factors like reduced concentration of glutathione which scavenges formaldehyde as S-hydroxymethylglutathione might be responsible for the lack of complementation. Of note, in the search for nutritional factors that can trigger selection for *E. coli* ST131 we discovered that pectin can serve as a carbon source for the ST131 strains and other *E. coli* isolates (Supplementary Figure 11). This feature might contribute to establishment of *E. coli* in the gastrointestinal tract and its subsequent dissemination. The role of other genome alterations still needs to be investigated.

Although an *E. coli* ST131 isolate has not been recovered upon the onset of the wound infection, we speculate that an *E. coli* ST131 isolate had been initially acquired from the environment, but remained undetected due to low numbers. ST131 strains had subsequently been selected during long term antimicrobial treatment and, potentially, due to an advantage to persist in the wound environment. If so, the ST131 clone isolate persisted despite of a less extended antimicrobial resistance profile. *E. coli* ST131 clone isolates are known for their ability to persist in the host and the environment ^7, 65^. The infection spectrum of ST131 isolates is wide and ST131 strains have been reported to occasionally even cause keratitis, and meningitis ^66, 67^. Further more an anecdotical observation reported a chronic tissue infection with an *E. coli* ST131 strain from an environmental source with horizontal transfer of plasmid from a co-infecting *Morganella* strain ^68^. Thus, environmental acquisition of ST131 by an open wound leading to a persistent infection does occur under certain circumstances. The yet-to-be-discovered molecular mechanisms which provide the *E. coli* ST131 clonal members with an exceptional capability to persist might also aid its long-term survival on foreign bodies and sequester in host tissue.

One alteration common to all ST131 isolates including the Bone strains is the truncated cyclic di-GMP dependent phosphodiesterase YcgG which consists only of the catalytic EAL domain deleted by its N-terminal signaling domain with still, although altered, functionality ^57, 69^ equally as substantial amino acid substitutions in other cyclic di-GMP turnover proteins compared to E. coli reference strain *E. coli* K-12 MG1655. Gene products with amino acid substitutions and/or extensions and truncations can have acquired an altered or novel functionality and/or expression pattern which might aid persistence of ST131 and Bone isolates in tissue ^70^.

Most surprisingly, however, was the delayed development or lack of biofilm formation in established assays such as formation of the rdar colony morphology type and biofilm formation on the polystyrene surface, respectively^71^ indicating that unconventional mechanisms contribute to the persistence of ST131 Bone isolates in the wound.

## MATERIALS AND METHODS

### Bacterial strains

Bacterial strains and their characteristics are summarized in Supplementary Table 1. Investigated *E. coli* isolates were Bone1A and B (isolated 01/01/2005), Bone2 (isolated 01/01/2005), Bone 4 (isolated 09/06/2013), Bone8 (07/07/2015), Bone7 (24/10/2015) and *E. faecalis* (24/10/2015). Reference *E. coli* strains were the laboratory strain *E. coli* K-12 MG1655, the commensal isolate TOB1 and the ST131 strain EP436. Biofilm reference strains were *Salmonella typhimurium* UMR1 (regulated biofilm formation) and MAE52 (constitutive high biofilm formation). Strains were grown on agar plates containing medium: LB without salt, blood, Mueller Hinton, Mueller Hinton fastidious and CLED and in Mueller Hinton and Bolton broth base. Antibiotics were used at the following concentrations; ampicillin Amp, 100 µg/ml; chloramphenicol Cm 25 µg/ml; tetracycline Tc 15 µg/ml.

### Phenotypic characterisation

Resistance against antibiotics was experimentally determined according to European Committee on Antimicrobial Susceptibility Testing (EUCAST) protocols and standards.

Assessment of the synthesis of the extracellular matrix components, the exopolysaccharide cellulose and amyloid curli fimbriae displaying cyclic di-GMP dependent rdar biofilm formation in the agar plate model was performed as described ^72, 73^. Congo red agar plates without or with the addition of 11 mM glucose were incubated at 28 or 37 °C for up to 72 h. TOB1_rdar28/37_, MAE52_rdar28/37_ and UMR1_rdar28_ serves as positive controls, while MAE777_saw28/37_ served as negative control. Colony morphology was visually documented using a conventional mobile phone. Of note, Congo Red agar plate growth can indicate membrane integrity, with microbes showing impaired cell envelope integrity displaying extensive dye uptake and subsequently stallation of growth.

Biofilm formation on an abiotic surface in the 96-well-plate format was performed to assess rdar biofilm formation and type 1 fimbriae dependent biofilm formation. Inoccula were incubated in 200 µl LB without salt broth at 28 °C for 48 h and LB broth at 37 °C for 24 h, respectively. Culture were assessed for pellicle formation and cell aggregation as an alternative biofilm format before careful aspiration of the medium. Attached biofilm cells were stained with a 0.2% crystal violet solution for 10 min and subsequently washed with destilled water until the washes did not contain any unbound dye (conventionally three times washing).

Flagella dependent swimming motility of isolates was assessed in Bolton broth medium with 0.25% agar ^2^. A colony of the isolate or three µl of a suspension of 5 OD was stabbed into the motility agar and the plate incubated at 37 °C with observation of motility at each hour until 9 hours and afterwards at 16 and 24 hours for slowly swimming isolates. *E. coli* MG1655 served as a positive control and *S. typhimurium* MAE108 (Δ*fliC* Δ*flgB*) as a negative control.

CLED agar was used to assess lactose fermentation of isolates.

Growth was either measured in 96 well plates format at 37°C without or with shaking (200 µl medium; *SpectraMax i3x*). or in Erlenmeyer flasks filled with 60 and 10% medium to provide microaerophilic and aerobic conditions, respectively ^74^. LB medium, Bolton broth, Bolton broth with hemin or Bolton broth with 5% lysed red blood cells (RBC) was the growth medium.

Screen for metabolic capability including use of carbon and energy sources was performed using the Biolog GEN III microplate test panel in three technical replicates. The use of pectin as carbon source was verified in an individual growth experiment.

### Preparation of lysed red blood cells

Lysed red blood cells were prepared by centrifugation of whole human blood at 500xg for 10 min at 4° C. The plasma was aspirated and the erythrocyte pellet washed with phosphate-buffered saline (PBS). After a second round of centrifugation and washing, the red blood cells were frozen at -20° C. A stock solution of hemin (50 mg/ml) was prepared in 0.05 M NaOH.

### Isolation of high molecular weight genomic DNA

High molecular weight genomic DNA was isolated by three different procedures; with the QIAGEN Genomic-tip 500/G using the provided buffers and following the instructions of the manufacturer with modifications, using the QIAGEN protocol using CHCl_3_ extraction instead of silica column purification and using phenol/CHCl_3_/isoamylalcohol (25/24/1) extraction. Purity and integrity of the extracted DNA was analyzed by spectral analysis measuring the 260/280 and 230/260 nm absorbance ratio by NanoDrop and by running the genomic DNA on a 0.8% agarose gel in 1x TAE buffer, respectively.

### Pulsed-field gel electrophoresis

Genomic DNA from bacterial isolates was prepared encapsulated in agarose plugs as described ^75^ and subsequently subjected to restriction digest by XbaI, BlnI and I-CeuI. Fragments were separated using a CHEF-DRII PFGE apparatus from Biorad in a 1% agarose gel in 0.5× Tris-Borate-EDTA at 6 V/cm and 14°C using switch times from 2 to 54 sec for 20 h (XbaI).

### Whole genome sequencing

Whole genome sequencing was performed with the PacBio RS II system (Pacific Biosciences at NGI Uppsala, Science For Life Laboratory [SciLifeLab], Uppsala, Sweden). The assembly was performed on SMRT portal version 2.3, with HGAP3 default settings. All strains were also sequenced on the Illumina MiSeq version 3 platform with read length up to 2 x 300 bp (NGI Stockholm, SciLifeLab, Solna, Sweden). The first genome sequence in 2018 did not provide sequences of acceptable quality for the ST131 Bone isolates. Subsequently, genomic DNA from each strain was sequenced at least two times isolated by the different experimental approaches. Sequencing results were compared with ambiguous sequence stretches resolved by PCR analysis, restriction digest and, if required, Sanger sequencing. Primers are listed in Supplementary Table 5.

### Cloning of genes

The *hemB* gene including its Shine-Dalgarno sequence was cloned by ligating XbaI/SphI restriction enzyme digested fragment and pBAD28 vector in vitro with subsequent transformation into chemocompetent *E. coli* TOP10 cells and selection of ampicillin resistant colonies.

All other genes were cloned by in vivo cloning ^76, 77^ with defined insertions in the vector mimicking conventional restriction site cloning. Shortly, open reading frames (with (pBAD28) or without (pSRKTc) Shine-Dalgarno sequences) were amplified with primers containing 15 bps complementary to the vector sequence at the 5’ end with XbaI/SphI and NdeI/NehI restriction sites at the immediate 3’ end. pBAD28 and pSRKTc vectors were amplified with primers containing the relevant restriction sites at their 5’ end. The two fragments were transformed into chemocompetent *E. coli* TOP10 cells and antimicrobial resistant colonies selected.

Colonies were screened by PCR for the insert, recombinant plasmids isolated and sequence integrity of the insert assessed by Sanger sequencing. Vectors and constructed plasmids are summarized in Supplementary Table 6 and primers in Supplementary Table 5.

### Deletion of chromosomal genes

The *hemB* gene was deleted by One-Step gene deletion according to established methodology ^78^ with the following modifications. To match with the antimicrobial resistance profile of the ampicillin resistant Bone4 strain, the gene cassette mediating ampicillin resistance in the λRed recombinase bearing pKD46 was replaced by *tetRA* mediating tetracycline resistance by in vivo cloning with the subsequent selection for tetracycline resistant colonies to obtain pKD66.

### Bioinformatic analysis

Chromosomes and plasmid sequences were compared with YASS ^79^ and gVISTA ^80^ to identify breakpoint of the structural genome variants and larger deletions and insertions. The genomes were annotated with RASTtk, Proksee and in MicroScope ^81-83^ for preliminary analysis. Illumina sequence reads for Bone 4, Bone 8 and Bone 7 were mapped onto the PacBio RII created assembly in UGENE ^84^. Assemblies of genomes of the same and different strain were analyzed using YASS ^79^, mVISTA ^85^, and SnapGene. Genome sequences were subsequently submitted to the NCBI database under the project number PRJNA634765 and annotated with the NCBI prokaryotic genome annotation pipeline (PGAP; ^86^). BLAST was used to assess protein homologies and nucleotide sequences ^87^, ClustalX2 ^88^ and UniProt was used to countercheck annotations ^89^.

Determination of the MLST types was done with http://bacdb.org/BacWGSTdb/Tools_results_multiple.php ^39^ assessing 7-gene MLST (*adk*, *fumC*, *gyrB*, *icd*, *mdh*, *purA* and *recA*). Virulence genes were preliminary identified by VirulenceFinder 2.0; search with standard parameters, % identity >90% and minimum length 60% (https://cge.cbs.dtu.dk/services/VirulenceFinder/). Resistance genes were preliminary detected by ResFinder 4.1 (https://cge.cbs.dtu.dk/services/ResFinder) and the Comprehensive Antibiotic Resistant Database (CARD; https://card.mcmaster.ca/) to assess acquired and intrinsic resistance genes. All identified loci were manually verified and curated. Enterobacteriales Origin of replication has been identified by PlasmidFinder 2.0.1 with standard parameters of 95% identity and 60% minimum coverage (https://cge.cbs.dtu.dk/services/PlasmidFinder/) ^90^. FimH alleles were determined by FimFinder 1.0 (https://cge.cbs.dtu.dk/services-/FimTyper) and serotype was determined by SerotypeFinder all at the Centre of Genomic Epidemiology website (http://www.genomicepidemiology.org/). pMLST was used to assess the plasmid type ^91^. Phages were identified by PHASTER and PHASTEST ^92, 93^.

### Phylogenetic analysis of ST131 genomes

To perform genome-based phylogenetic analysis by core genomes, FASTA genomic sequence files of the representative ST131 from different clades were downloaded from the NCBI, ENA and EnteroBase websites considering the completeness of the individual genomes. To ensure equal analysis treatment of the different genomes, reannotation of the genomes was performed using the annotation package Prokka 1.14.5 ^94^. Subsequently, the core genomes from the target data set were calculated by Roary software 3.11.2 with a threshold of 90% identity ^95^. Alignments from the conserved genes were concatenated to calculate a maximum-likelihood genome-based phylogenetic tree using PhyML 3.0 of the SeaView software 5.0.4 ^96^. If not otherwise indicated, the respective reference genomes were used.

### Visualisation tools

Compariative analysis of genome and plasmid sequences was analysed and visualized with EasyFig ^97^. Circoletto and Proksee was used to visualize, annotate and compare plasmid sequences with annotation performed within the programmes ^83, 98^. MEGA 7.0 was applied to visualize the phylogenetic tree of ST131 genomes ^99^.

## Acknowledgement

The authors thank Erik Holmqvist, Uppsala University for assessment of the integrity of ProQ by Western blotting at an early stage of this project. NOD and MSG were supported by RSF grant 24-14-00276. SKG was supported by the Intramural Research Program of the National Library of Institutes of Health (NIH). The contributions of the NIH authors are considered Works of the United States Government. The findings and conclusions presented in this paper are those of the authors and do not necessarily reflect the views of the NIH or the U.S. Department of Health and Human Services. UR has been supported by the Petrus and Augusta Hedlund Foundation, ALF (Swedish National Agreement on Clinical Research and Training) and the Swedish Research Council for Natioanl Sciences and Engineering.

The authors would like to acknowledge support of the National Genomics Infrastructure (NGI) /Uppsala Genome Center and UPPMAX for providing assistance in massive parallel sequencing and computational infrastructure. Work performed at NGI/Uppsala Genome Center has been funded by RFI/VR and Science for Life Laboratory, Sweden.

---

**Supplementary TABLE S1** Bacterial strains isolated and used in this study

**Supplementary TABLE S2** Basic genomic and plasmid characteristics of Escherichia coli Bone isolates compared to ST131 clade C1 and C2 reference strains.

**Supplementary TABLE S3** Amino acid alterations in the c-di-GMP metabolizing proteins of Bone isolates and ST131 reference strains compared to the respective reference protein sequences in *E. coli* MG1655.

**Supplementary TABLE S4**

A Insertions and deletions of Bone4 compared to reference ST131 clade C1 isolate AR_0058.

B Single nucleotide polymorphisms and insertions <20 bps of Bone4 compared to reference ST131 clade C1 isolate AR_0058.

C Insertions and deletions between Bone4 and Bone5, Bone8 and Bone7.

D Single nucleotide polymorphisms between Bone4 and Bone5, Bone8 and Bone7.

E Inverting regions within Bone4, Bone5, Bone8 and Bone7 isolates.

F Gene products encoded by chromosomal insertions/deletions between ST131 Bone and reference AR_0058 reference strain.

**Supplementary TABLE S5** Primers used in this study.

**Supplementary TABLE S6** Plasmids used in this study.

