## Supplemental Information Methods and Figures for "Evolution towards small colony variants of pandemic multidrug resistant ST131 *Escherichia coli* isolates from a 10-year wound infection"

### Materials and Methods

#### Quality assessment of genome assemblies after whole genome sequencing

Whole genome sequencing by PacBio RS II system (Pacific Biosciences at the National Core Facility NGI Uppsala, Science For Life Laboratory [SciLifeLab], Uppsala, Sweden) has been performed with genomic DNA isolated by three different methods. Q=modified Qiagen protocol; QC=modified Qiagen protocol with final DNA purification by $CHCl_3$ instead of the silica-column; P=conventional phenol/$CHCl_3$/isoamylalcohol 25:24:1 extraction. The following DNA isolations were snd for PacBio sequencing (in bold, reference genomic sequence: Bone4_Q, Bone3_2QC; Bone5P, Bone5Q, Bone5_QC; Bone7P, Bone7Q, Bone7_QC; Bone8_1QC, Bone8_2QC, Bone8P, Bone8Q. For one strain, Chromosome and plasmid assemblies were compared and inconsistencies assessed by PCR and, if required, Sanger sequencing. Two major regions of inconsistencies in the assembly were based on inversions of DNA fragments in phages.

Additional primers used to assess inconsistencies (deletion in the Bone4Q assembly) were: B4Q_R1_330939: CTGTAGACAGCAGCTCCACACC and B4Q_R1_332110: TGTCTCAGTTCCAGTGTGGCTG

#### Assessment of recombination events in ribosomal RNA operons

Ribosomal RNA operons in ST131 reference strains are found at the same distance, location and direction with reference to the origin of replication as in the *E. coli* K-12 MG1655 reference strain (REF Maeda, Ishihama Plos One 2015). While *rrnC*, *rrnA*, *rrnB*, *rrnE* and *rrnH* are found clock-wise arranged, *rrnD* and *rrnG* are found counterThe clockwise with with transcription in parallel with proceeding of DNA replication. In Bone4, recombination occurred between the *rrnH* and *rrnD* operons with the precise inversion breakpoints at [16S rRNA, tRNA-Ile, tRNA-Ala, 23S rRNA, 5S rRNA, (inversion breakpoint), tRNA-Thr, 5S rRNA] and [16S rRNA, tRNA-Ile, tRNA-Ala, 23S rRNA, 5S rRNA, (inversion breakpoint), tRNA-Asp]. In Bone 7, recombination occurred between *rrnC* and *rrnD*, with the precise inversion breakpoints at [16S rRNA, tRNA-Glu, 23S rRNA, 5S rRNA, (inversion breakpoint), tRNA-Thr, 5S RNA] and [16S rRNA, tRNA-Ile, tRNA-Ala, 23S rRNA, 5S rRNA, (inversion breakpoint), tRNA-Asp, tRNA-Trp].

### Supplementary Figures

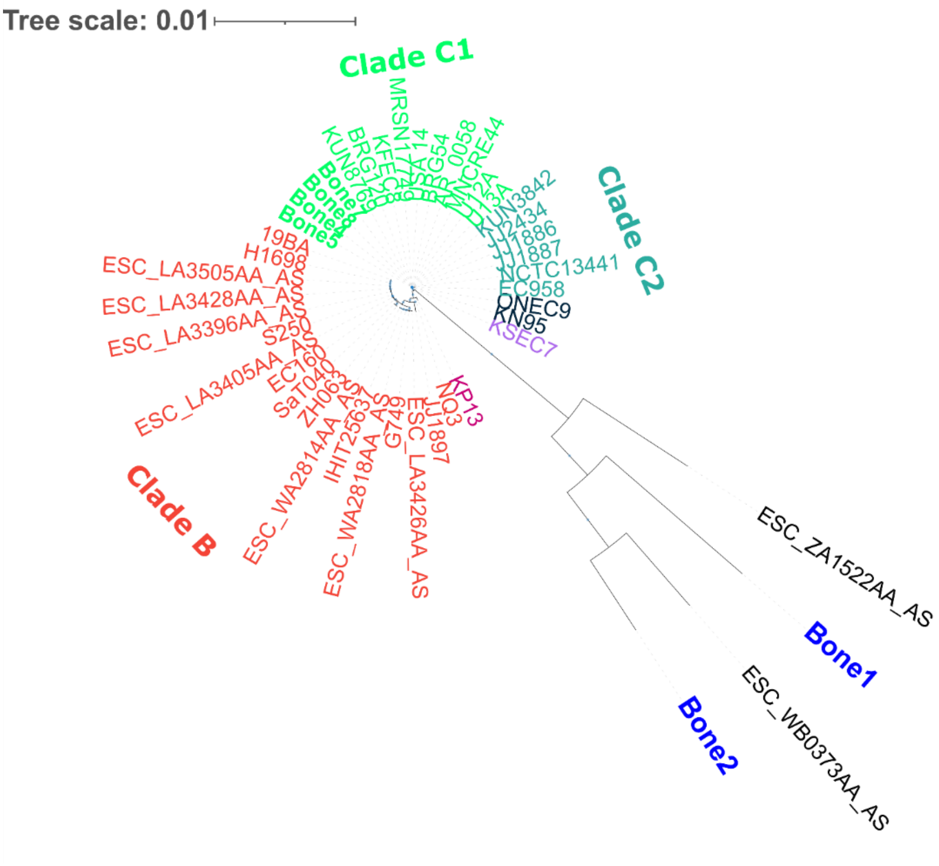

**Supplementary FIG 1** Phylogenetic classification of the ST131 Bone4,5,7 and 8 isolates versus Bone1 and Bone2 isolates. 37 ST131 *E. coli* strains had been selected to represent the clinically most relevant ST131 clades B, C1, C2 and minor clade B0 and C0. Phylogenetic tree based on the core genome alignment.

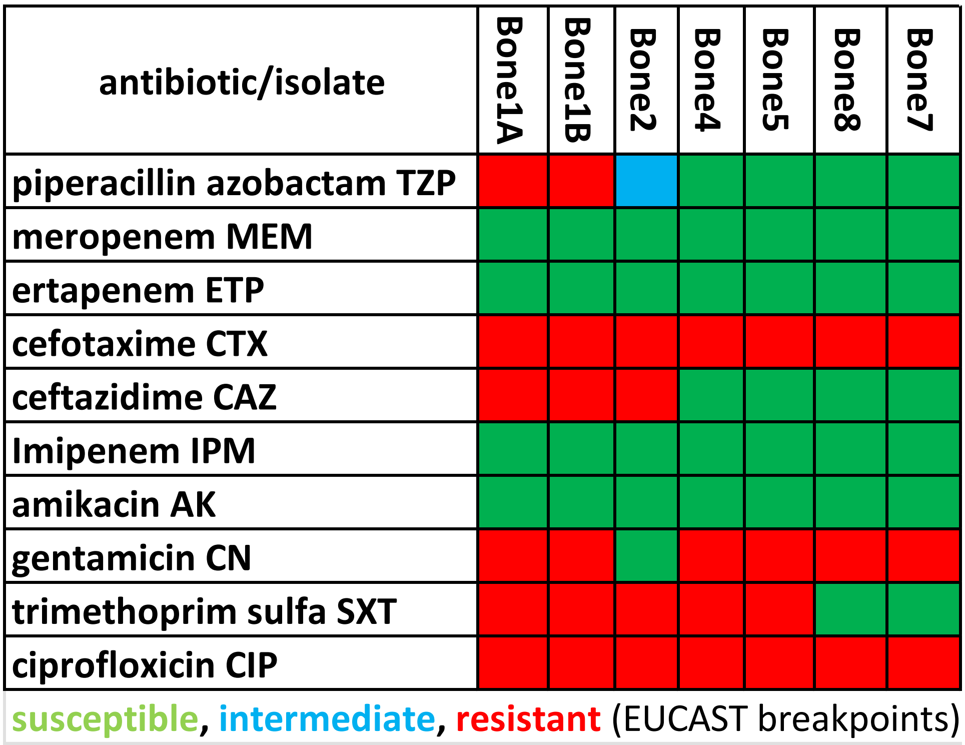

**Supplementary FIG2** Heatmaps of experimentally determined resistance profiles and identified chromosomal target mutations and chromosomal and plasmid encoded antibiotic resistance determinants.

A Experimentally determined resistance profiles for representatives of clinically most relevant antibiotic classes.

| chromosomally encoded antibiotic resistance determinants | Bone1A | Bone1B | Bone2 | Bone4 | Bone5 | Bone8 | Bone7 | resistance |
|---|---|---|---|---|---|---|---|---|
| **gene cassettes** | | | | | | | | |
| *blaCTX-M-14* | green | green | green | red | red | red | red | third generation cephalosporines |
| *blaCTX-M-15* | green | green | red | green | green | green | green | third generation cephalosporines |
| class C EC-18 | red | red | green | green | green | green | green | beta-lactam antibiotics |
| class C EC-8 | green | green | red | green | green | green | green | beta-lactam antibiotics |
| *terC1, terC2* | red | red | red | red | red | red | red | tellurite resistance |
| SAT-2 | red | red | green | green | green | green | green | streptothricin, nourseothricin |
| *aadA1* | red | red | green | green | green | green | green | streptomycin, spectinomycin |
| *tet(B)* | red | red | red | green | green | green | green | tetracycline, doxycycline |
| *dfrA1* | red | red | green | green | green | green | green | trimethoprime |
| *sitABCD* | green | green | red | red | red | red | red | hydrogen peroxide (combinatory) |
| **target mutations fluoroquinolones** | | | | | | | | |
| GyrA | red | red | red | red | red | red | red | fluoroquinolone |
| GyrB | green | green | red | red | red | red | red | fluoroquinolone |
| ParC | red | red | red | red | red | red | red | fluoroquinolone |
| ParE | red | red | red | green | red | green | green | fluoroquinolone |
| **target mutations other antibiotics** | | | | | | | | |
| HemA/HemB/HemL | green | green | green | green | green | dark red | dark red | protamine sulphate, gentamicin |
| EF-Tu | red | red | red | red | red | red | red | pulvomycin/elfamycin |
| GlpT | red | red | red | red | red | red | red | fosfomycin |
| CyaA | green | green | red | green | green | green | green | fosfomycin |
| UhpT | red | red | red | red | red | red | red | fosfomycin |
| PtsI | green | green | green | red | orange | green | green | fosfomycin |
| MarR | red | red | red | red | red | red | red | MDR |
| SoxR | red | red | red | red | red | red | red | MDR |
| SoxS | red | red | red | red | red | red | red | MDR |
| PBP3 | red | red | red | red | red | red | red | chephalosporin |

| | |
|---|---|
| red | presence of resistance gene cassette or amino acid substitution in target |
| green | absence of resistance gene cassette or wild type target |
| dark red | deletion of *hemB* , but no observed protamine sulphate resistance |
| orange | uncertain substitution in target |

B Chromosomally encoded resistance determinants. Amino acid substitutions in target gene products (according to CARD and ResFinder 4.1) and presence of an antibiotic resistance cassette that mediate resistance are indicated in red. Of note, a streptothricin resistance cassette is encoded by the Bone1B, a CTX-M-15 β-lactamase by the Bone2 and a CTX-M-14 β-lactamase by the Bone4,5,8 and 7 chromosomes.

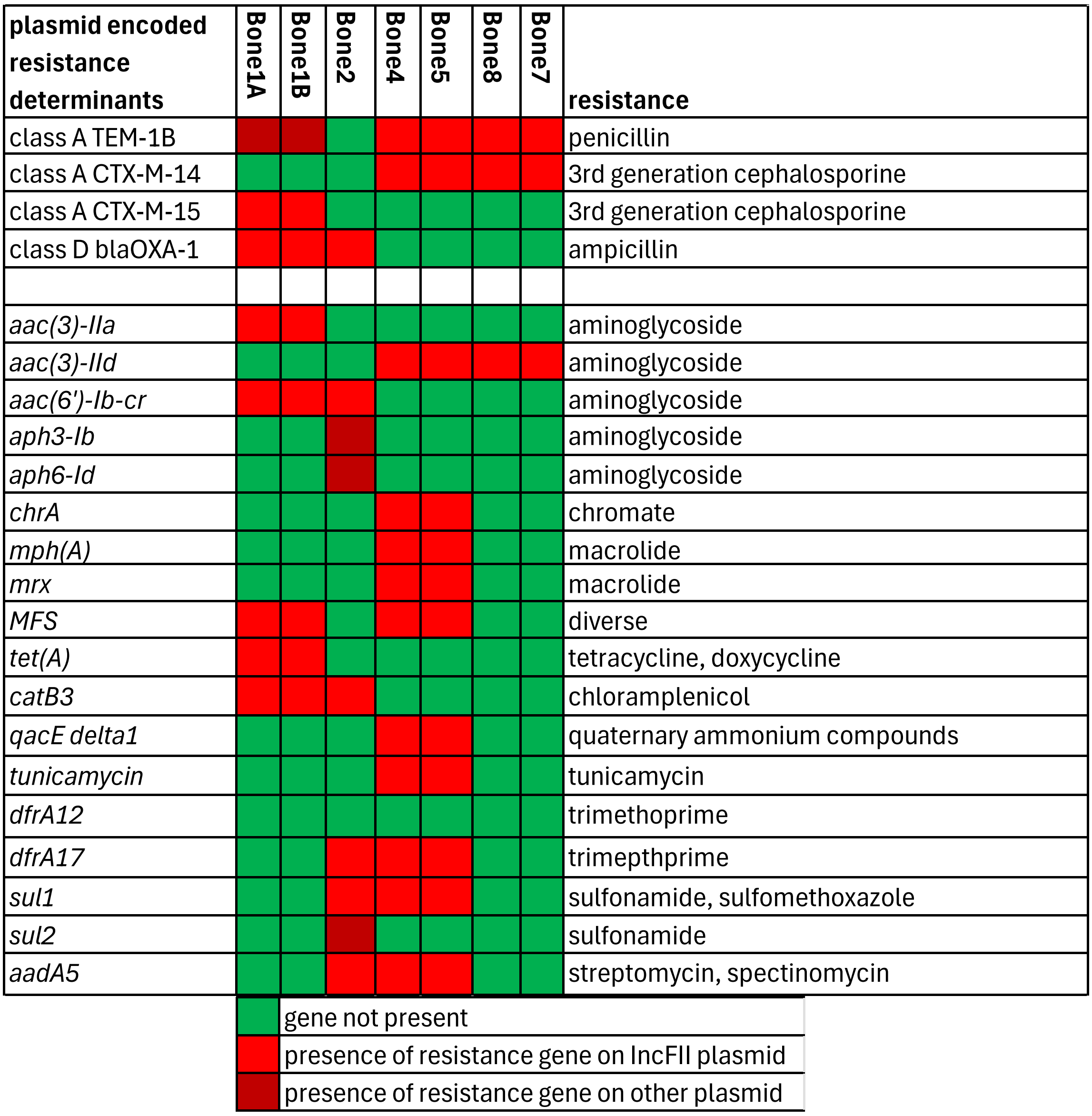

C Plasmid encoded resistance determinants. Resistance determinants: beta-lactamases: class A TEM-1B, class A CTX-M-14, class A CTX-M-15 and class D blaOXA-1. aminoglycoside resistance by *aac(3)-IIa* and *aac(3)-IId* encoding aminoglycoside-N(3)-acetyltransferases, *aph3-Ib* and *aph6-Id* encoding aminoglycoside 3'-phosphotransferase and *aadA5* encoding aminoglycoside nucleotidyltransferase; fluoroquinolone resistance by *aac(6')-Ib-cr* encoding aminoglycoside-(6)-N-acetyltransferase: chromate resistance by *chrA* encoding an efflux protein*;* macrolide resistance by *mph(A)* encoding macrolide 2´-phosphotransferase; *MFS* and *tet(A)* encoding multidrug transporters*;* chloramphenicol resistance by *catB3* encoding type B-3 chloramphenicol O-acetyltransferase*;* quaternary ammonium compound-resistance *qacE delta1; tunicamycin* resistance protein; trimethoprim resistance by *dfrA12* and *dfrA17* encoding dihydrofolate reductases; sulfonamide resistance *by sul1* and *sul2 e*ncoding dihydropteroate synthases. Of note, Bone1B and Bone2, but not the ST131 clade 1 strains Bone4, Bone8 and Bone7 harbor the class A CTX-M-15 beta-lactamase. Red, presence of the resistance cassette on the major IncFII plasmids; dark red, resistance cassette encoded by another plasmid; green, absence of the resistance cassette.

.

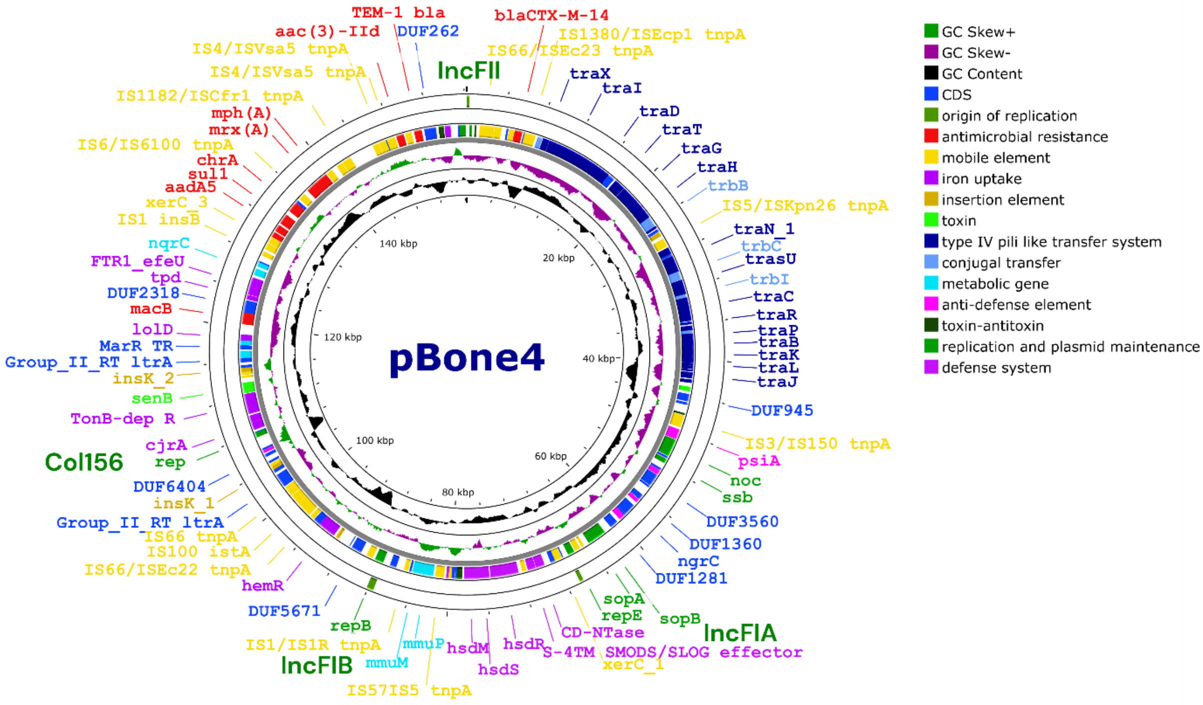

**Supplementary FIG 3** IncFII Plasmids in Bone isolates.

A Circular map of the 156.3 kbp large multireplicon IncFII plasmid pBone4. The sequence has been normalized to the IncFII origin of replication. An IncFIA, IncFIB and a Col156 origin of replication is also present as indicated by green bars in the third circle. Of note, antimicrobial resistance determinants cluster around the IncFII origin of replication. The 178 open reading frames, color coded according to functionality, are shown in the outermost circle. The G+C skew and G+C content is indicated as displayed. The plasmid map has been created with CG view [98] with programme embedded annotation of the open reading frames with Prokka and subsequent manual curation. Note that the annotation using Prokka[92] can differ from the annotation by the RAST server [99] and the prokaryotic pipeline of NCBI.

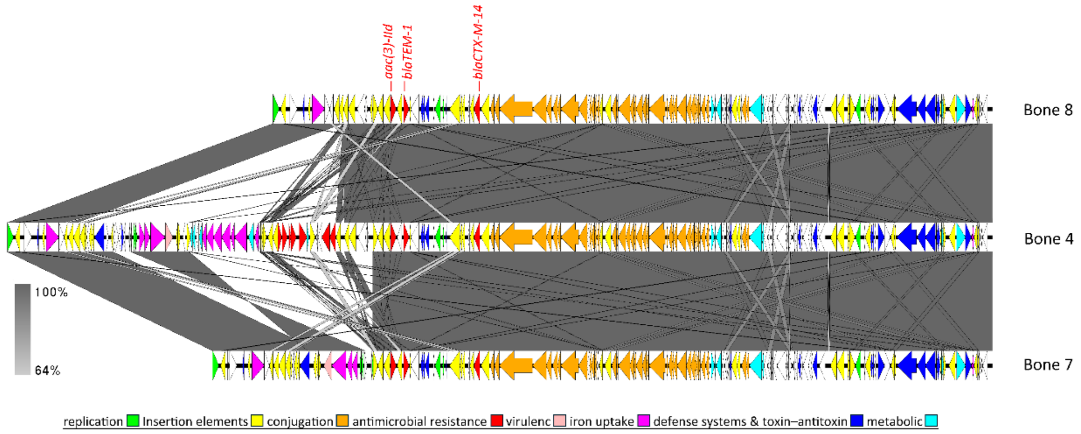

B Comparison of IncFII plasmids pBone4 with pBone8 and pBone7 plasmids. Sequences were normalized to the IncF1B origin of replication. pBone8 and pBone7 sequences are to a major part in synteny with pBone4 but lack approximately 25 kbp genetic information that encode also several antimicrobial resistance determinants. Sequence comparison and figure was created with EasyFig [95].

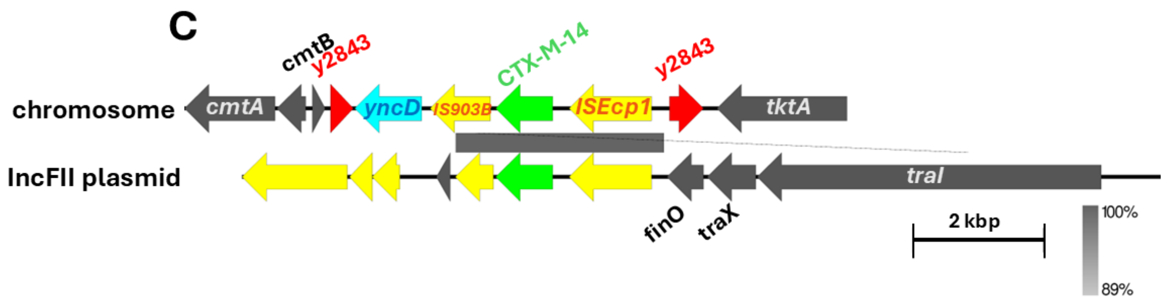

C Location of the CTX-M-14 β-lactamase gene on the IncFII plasmid and chromosome of ST131 Bone isolates. Translocation of the CTX-M-14 β-lactamase gene disrupts the y2843 protease.

D

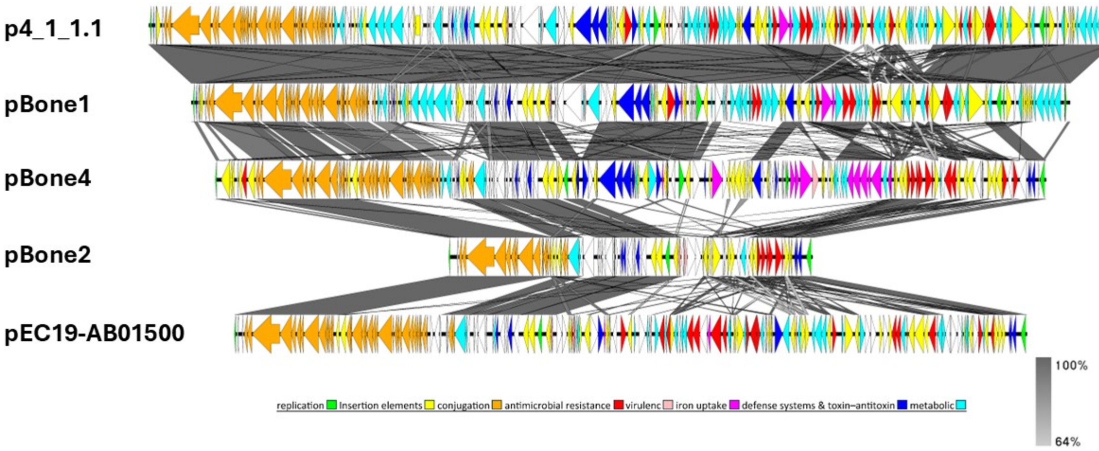

D Alignment of plasmid pBone1BIncFII, pBone2IncFII and pBone4 with plasmid most similar to pBone1BIncFII, p4_1_1 and pBone2IncFII, pEC19-AB01500 with localization of most homologous regions and major resistance determinants. Sequencees were normalized to the IncFII origin of replication. Most similar plasmids with largest query coverage had been identified by Blast search with pBone1BIncFII and pBone2IncFII nucleotide sequences as query. Annotated plasmids were aligned and visualized with EasyFig [95].

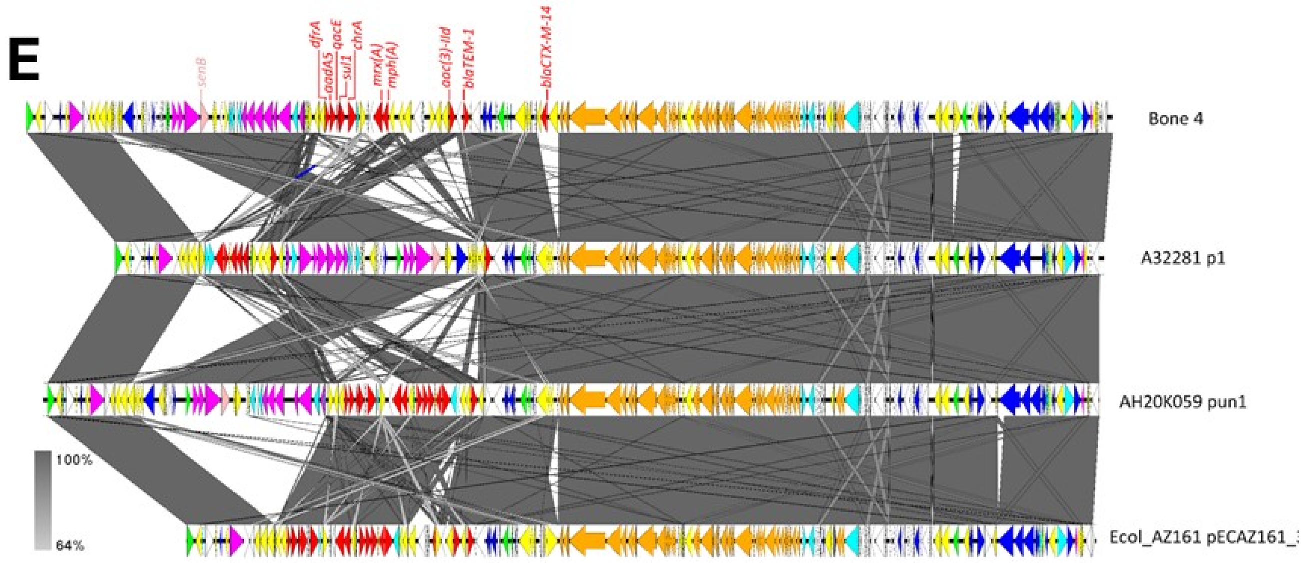

E Comparison of plasmid pBone4 with most similar plasmids as retrieved by Blast search with the nucleotide sequence of plasmid pBone4 as a query. With a conserved backbone, antimicrobial resistance determinants, virulence factors and iron uptake systems are variable.

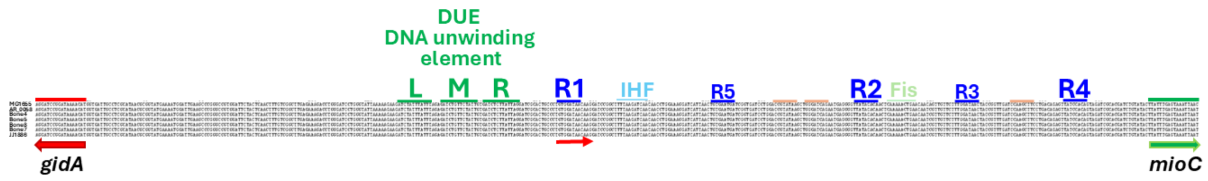

**Supplementary FIG 4** Normalisation of the ST131 chromosomes to the first base of the first high affinity DnaA binding site R1 at the origin of replication. The E. coli origin of replication is a 254 bp long region located between the *gidA* and the *mioC* genes. R1, 3 and 5 high affinity and R2 and 4 low affinity DnaA binding sites. Pink bars, other low affinity DnaA binding sites. L, M, R A/T-rich direct repeats of the DUE DNA unwinding element region, IHF and Fis protein binding sitesa re also indicated.

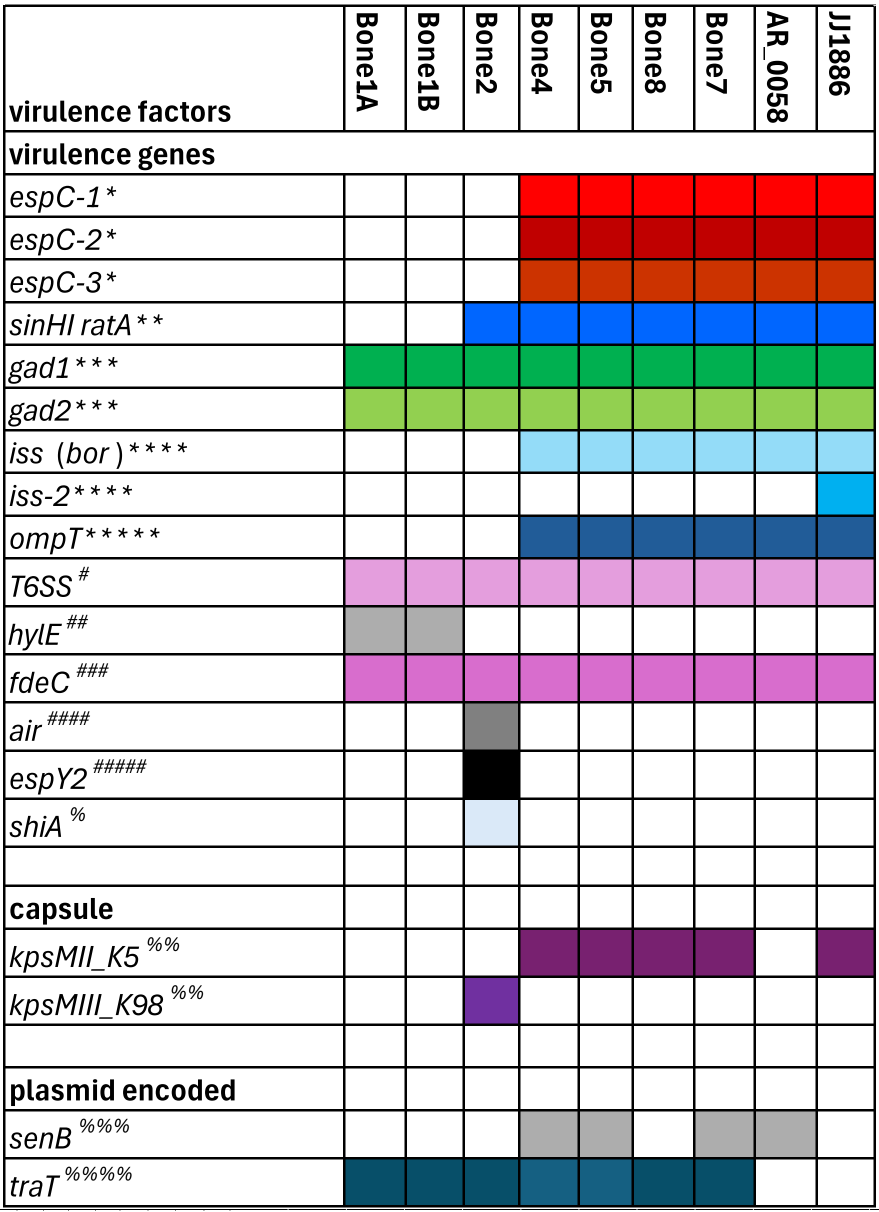

*SPATE, type5a secretion system serine protease autotransporter; ** invasin proteins; ***glutamate decarboxylase; ****prophage encoded increased serum resistance; *****outer membrane protease

#type 6 secretion system; ##avian specific haemolysin; ###inverse autotransporter intimin-like adhesin; ####intimin-like adhesin; #####type 3 secretion system effector

%immunomodulatory cytochrome c-like protein; %%capsule biosynthesis gene cluster; %%%enterotoxin; %%%%serum resistance

**Supplementary FIG 5** Comparative analysis of virulence factors, iron uptake systems and phages encoded by the Bone isolate genomes. AR_0058 and JJ1886 served as STM131 clade C1 and C2 references, respectively.

A Virulence factors encoded by the Bone isolate genomes. Virulence factors were search for by VirulenceFinder 2.0 and subsequently manually curated.

| iron uptake systems | Bone1A | Bone1B | Bone2 | Bone4 | Bone5 | Bone8 | Bone7 | AR_0058 | JJ1886 |
|---|---|---|---|---|---|---|---|---|---|
| **siderophore biosynthesis** | | | | | | | | | |
| enterobacin biosynthesis and transport *entDfepAfesybdZentFfepECGDentSfepBentCEBdhbAentH* | ■ | ■ | ■ | ■ | ■ | ■ | ■ | ■ | ■ |
| aerobactin *iutAiucA-D* | | | ■ | ■ | ■ | ■ | ■ | ■ | ■ |
| yersiniabactin biosynthesis | | | ■ | ■ | ■ | ■ | ■ | ■ | |
| **iron uptake and processing systems** | | | | | | | | | |
| Fe2+ uptake *efeUOB* | ■ | ■ | ■ | ■ | ■ | ■ | ■ | ■ | ■ |
| Fe2+ uptake *feoA-C* | ■ | ■ | ■ | ■ | ■ | ■ | ■ | ■ | ■ |
| Fe3+ dicitrate uptake *fecIRA-E* | | | ■ | ■ | ■ | ■ | ■ | ■ | |
| siderophore ferric coprogen receptor *fitA-E* | | | | ■ | ■ | ■ | ■ | ■ | ■ |
| TonB dependent receptor *cirA* | | | | ■ | ■ | ■ | ■ | ■ | ■ |
| TonB-dependent receptor *prrDCBhypomodDprrA* | | | ■ | ■ | ■ | ■ | ■ | ■ | ■ |
| ferrochrome binding protein *fhuA-D* | ■ | ■ | ■ | ■ | ■ | ■ | ■ | ■ | ■ |
| manganese ABC transporter *sitA-D* | | | ■ | ■ | ■ | ■ | ■ | ■ | ■ |
| TonB dependent ferrichrome receptor *hemR* -like+ *hmuS* | | | ■ | ■ | ■ | ■ | ■ | ■ | ■ |
| TonB-dependent catecholate siderophore receptor *fiu* | ■ | ■ | ■ | ■ | ■ | ■ | ■ | ■ | ■ |
| TonB-dependent receptor *iha* | | | | ■ | ■ | ■ | ■ | ■ | ■ |
| catecholate siderophore receptor *hma* | ■ | ■ | ■ | ■ | ■ | ■ | ■ | ■ | ■ |
| outer membrane hemin receptor *chuA* | | | | ■ | ■ | ■ | ■ | ■ | ■ |
| TonB-dependent receptor *fhuE* | ■ | ■ | ■ | ■ | ■ | ■ | ■ | ■ | |
| siderophore-iron reductase *fhuF* | ■ | ■ | ■ | ■ | ■ | ■ | ■ | ■ | ■ |
| iron export *fetAB* | ■ | ■ | ■ | ■ | ■ | ■ | ■ | ■ | ■ |
| Fha-like exoprotein in heme utilization and adhesion* | | | ■ | ■ | ■ | ■ | ■ | ■ | ■ |
| TonB dependent receptor *yncD* (*pqqU*)** | ■ | ■ | ■ | ■ | ■ | ■ | ■ | ■ | ■ |
| *, polymorph in Bone2; **most likely no iron receptor/transporter | | | | | | | | | |
| **on IncFII plasmid** | | | | | | | | | |
| $Fe^{2+}$ iron ABC transporter 03520 A-F | ■ | ■ | | ■ | ■ | ■ | ■ | | ■ |
| TonB dependent ferrichrome and hemi receptor | | | | ■ | ■ | ■ | ■ | ■ | |
| TonB family protein | | | | ■ | ■ | | ■ | | |
| TonB dependent receptor | | | | ■ | ■ | | ■ | | |

B Iron uptake systems encoded by the Bone isolate genomes. Iron uptake systems were partially detected by VirulenceFinder 2.0 and subsequently manually curated.

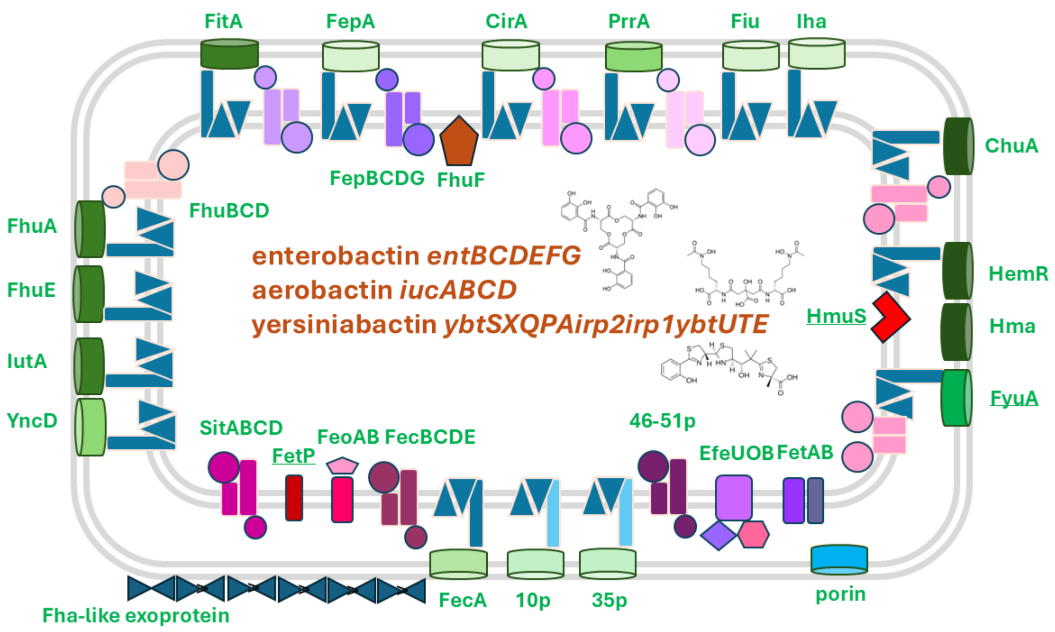

C Visualisation of iron uptake systems in ST131 Bone isolates. Siderophore structures from Wikipedia.

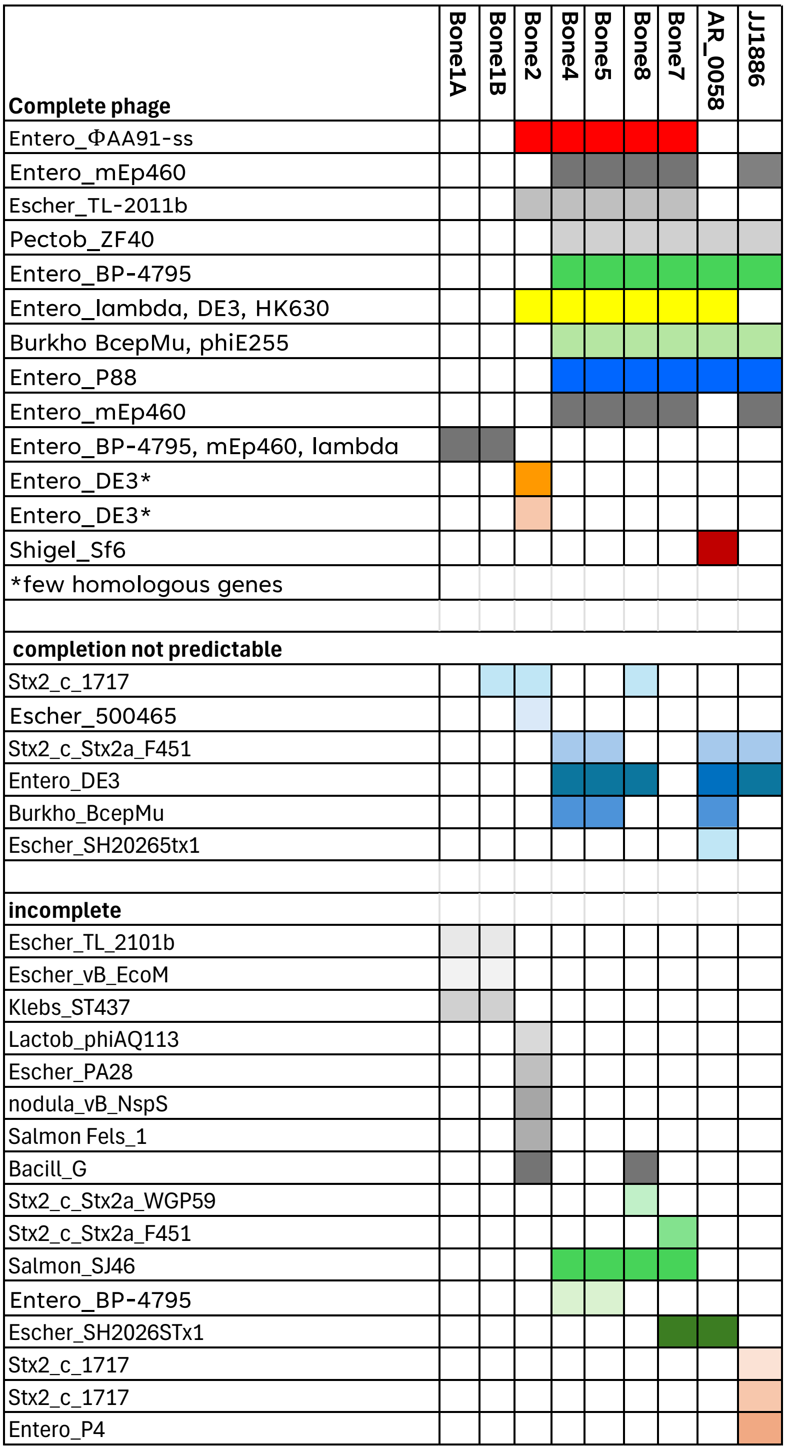

D Functional, conditionally functional and incomplete phages as recognized by Phaster and Phastest.

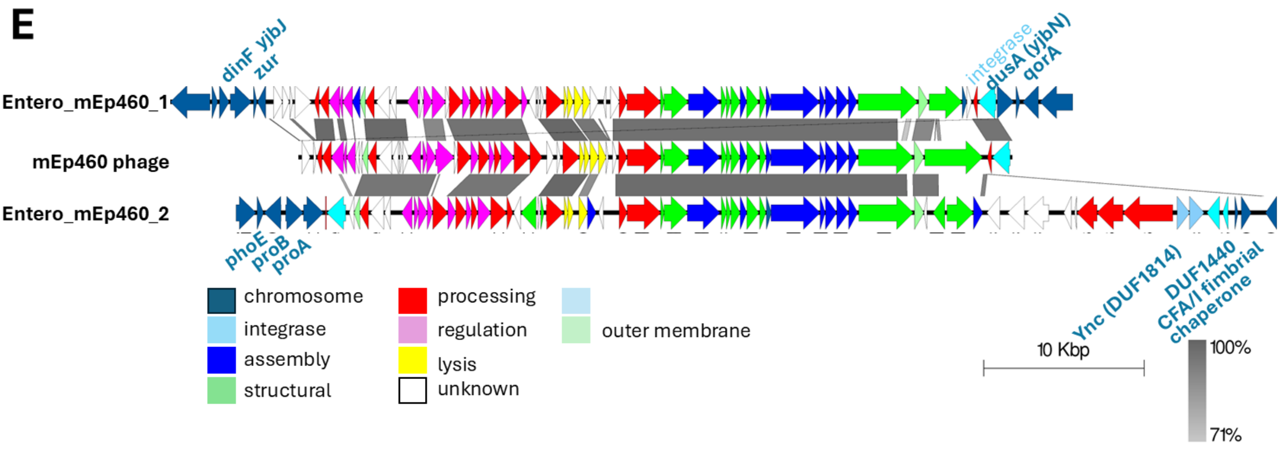

E Insertion site and genome organisation of the two mEP460 phages present in the ST131 Bone isolate chromosomes.

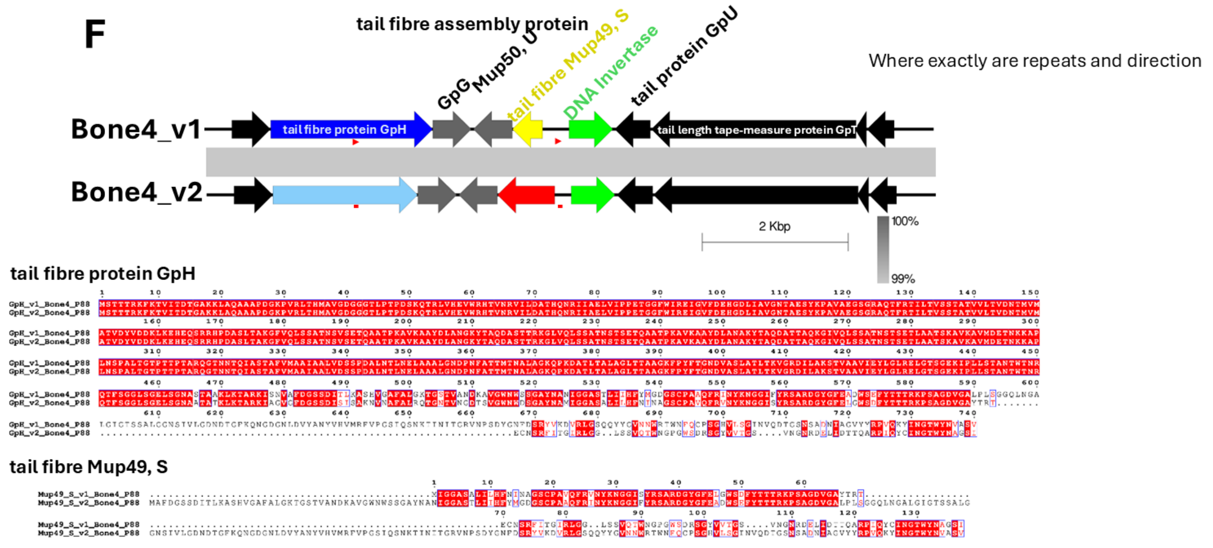

F Zoom in and alternative protein sequences for GpH and Mup49, S of the Entero_P88 phage created by the DNA inversion. Red arrow, direct repeat sequences of 70 bp.

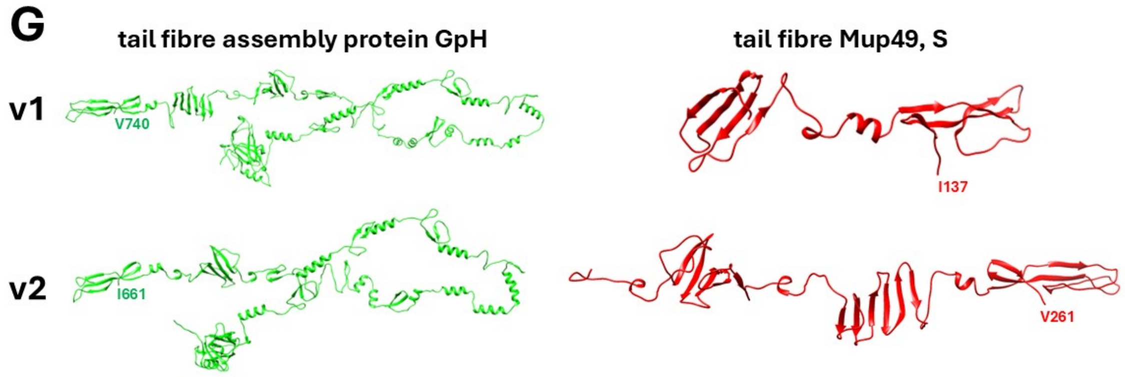

G AlphaFold 3 models of alternative proteins for tail fibre assembly protein GpH and teil fibre protein Mup49, S of the Entero_P88 phage.

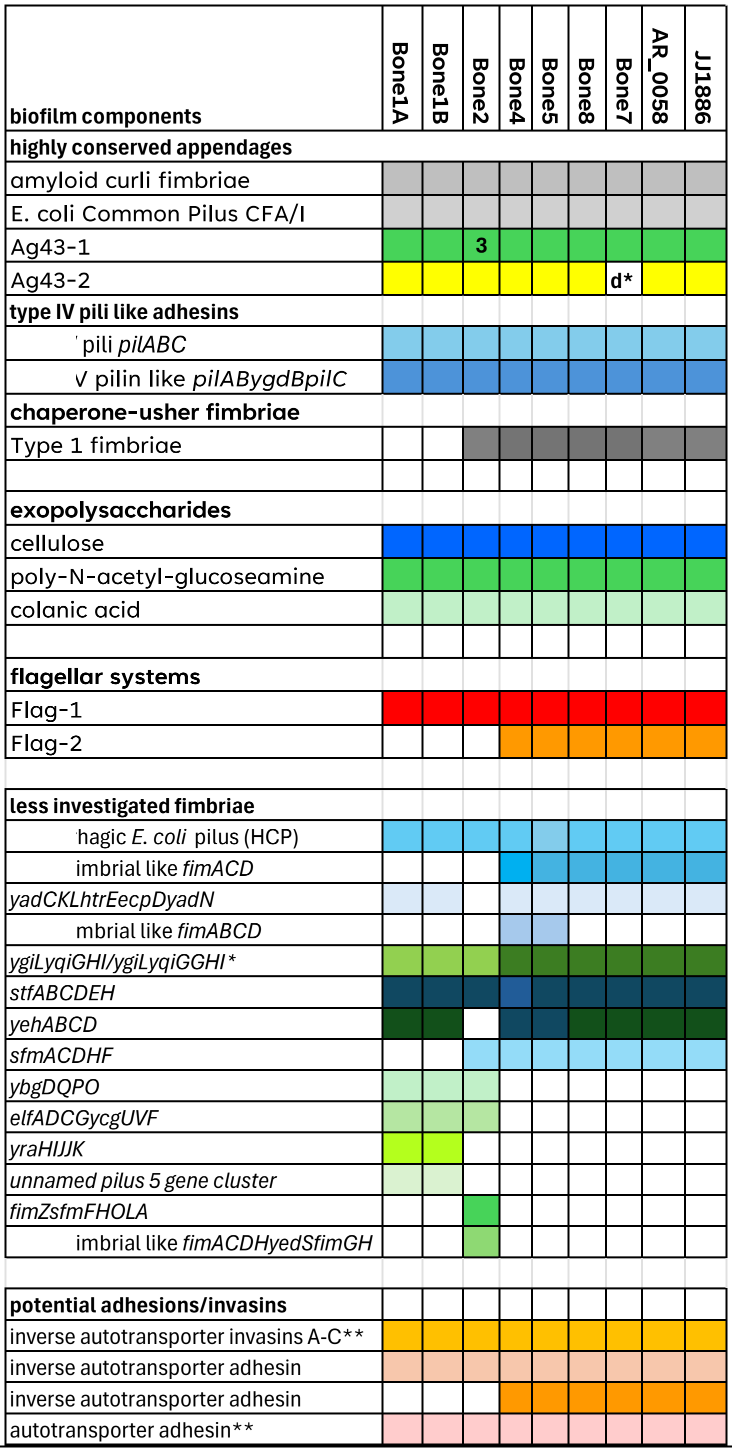

*d, disrupted, polymorphic YgiL 60% identity to ST131 protein;

**polymorphic, 80% identity in Bone1 and Bone1 and 2, respectively

**Supplementary FIG 6** Comparative analysis of occurrence and variability of biofilm components in Bone isolates. AR_0058 and JJ1886 served as STM131 clade C1 and C2 references, respectively.

A Comparative analysis of extracellular matrix components and adhesins reported or predicted to be involved in biofilm formation of *E. coli* isolates.

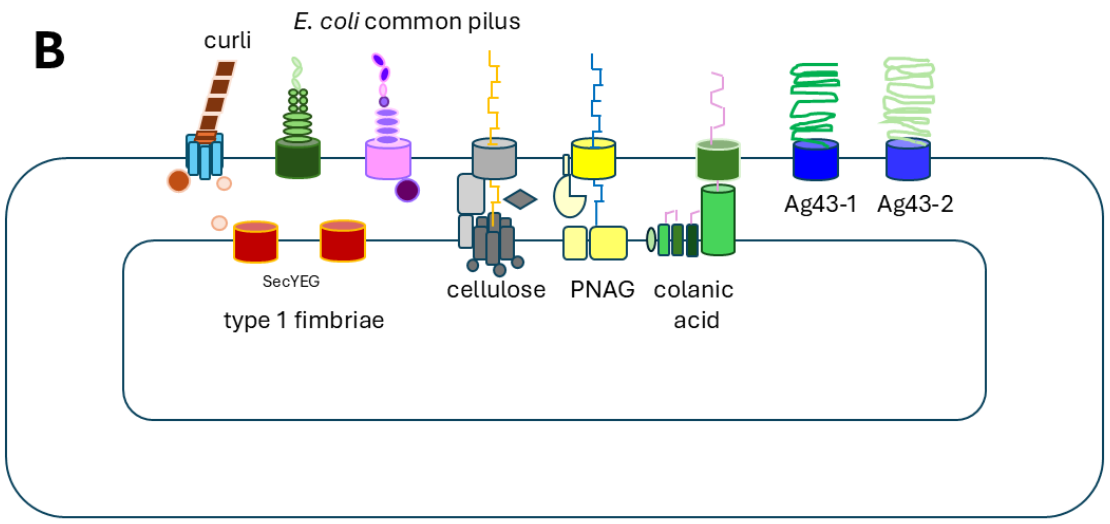

B Display of nanomachines involved in the synthesis of major biofilm extracellular matrix components of *E. coli.*

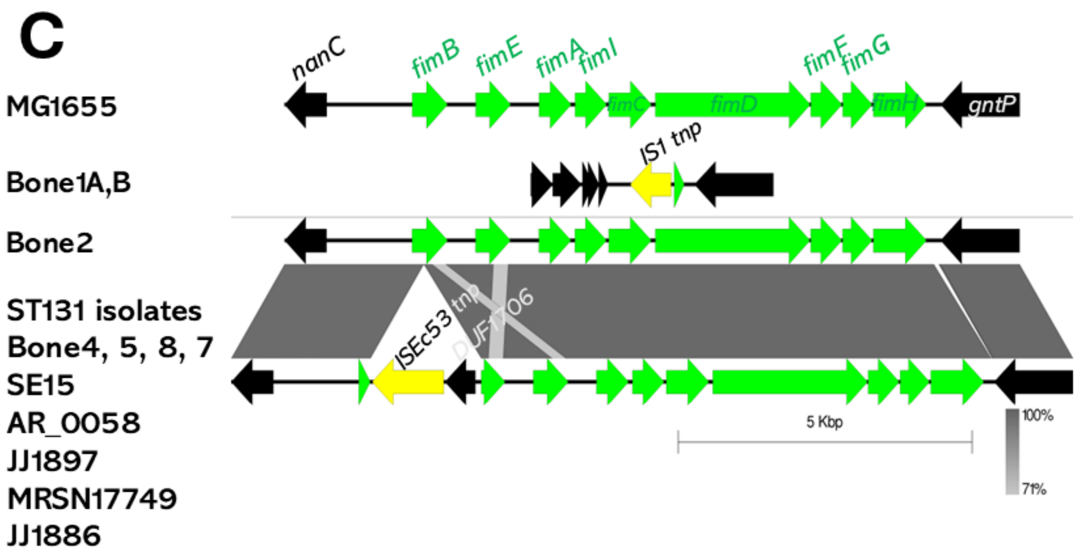

C Comparative analysis of the arrangement of type 1 fimbriae in Bone isolates. *E. coli* K-12 MG1655 served as well-documented reference.

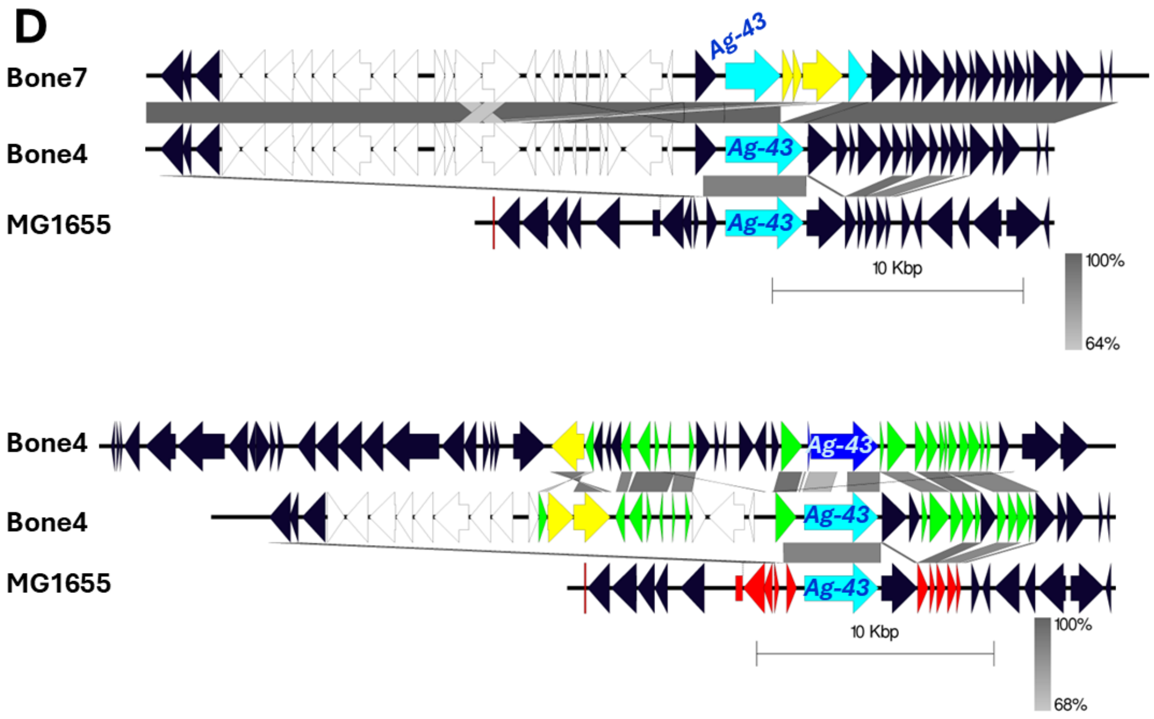

D Genomic context of the Ag43 adhesins in ST131 Bone isolates compared to *E. coli* K-12 MG1655 and displaying the inactivation event of Ag-43_1 in Bone7.

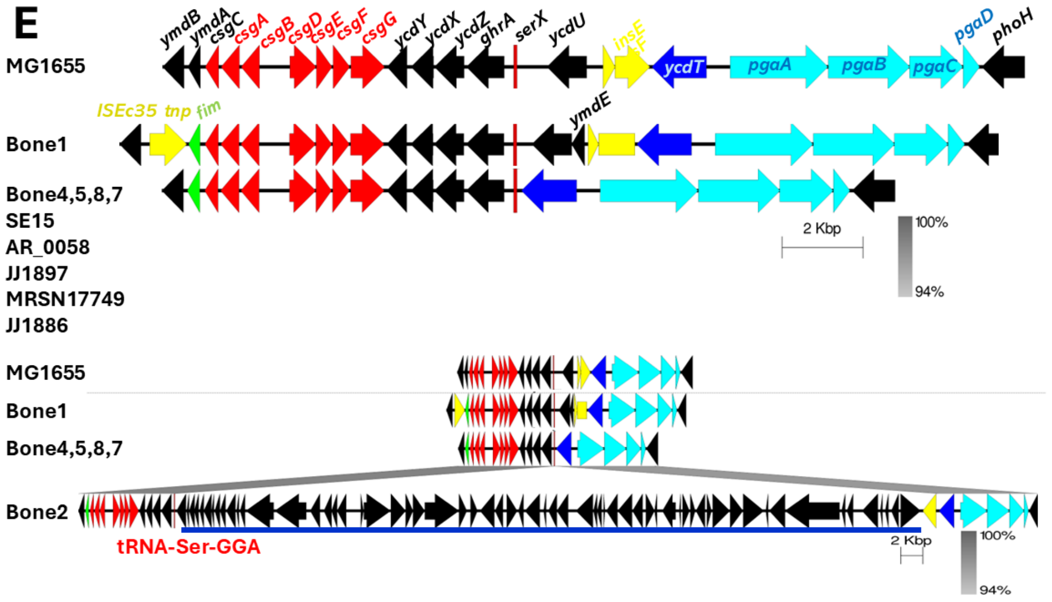

E Genomic context of the proteinaceous amyloid curli and exopolysaccharide poly-N-acetyl-glucoseamine operons in Bone isolates compared to well-documented reference *coli* K-12 MG1655. Bone2 has an insertion at the serine tRNA which codes for one copy of Ag43_1, restriction modification systems and metabolic gene products.

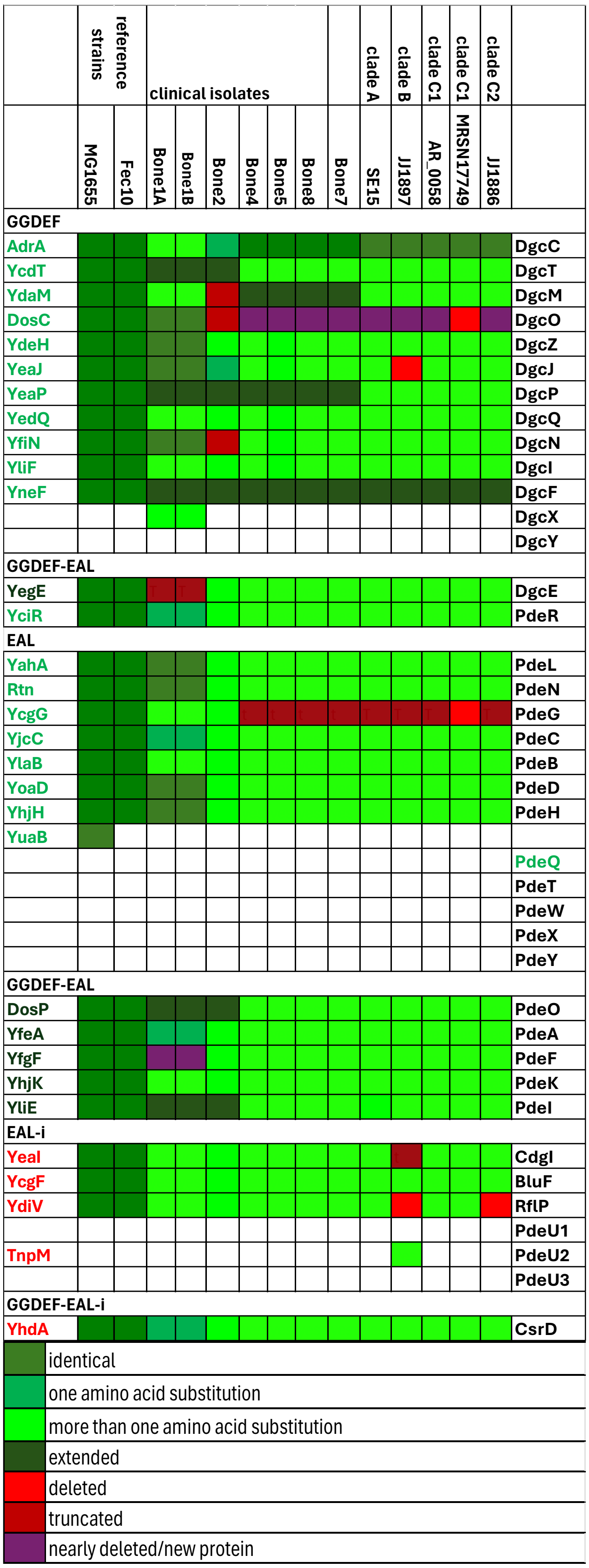

**Supplementary FIG 7** The cyclic di-GMP signaling network involved in the regulation of biofilm formation, motility and virulence.

A Heatmap with comparative analysis of cyclic di-GMP turnover proteins in Bone isolates compared to *E. coli* K-12 MG1655/Fec10 and ST131 clade reference strains assessed against all cyclic di-GMP turnover proteins previously observed in *E. coli.*

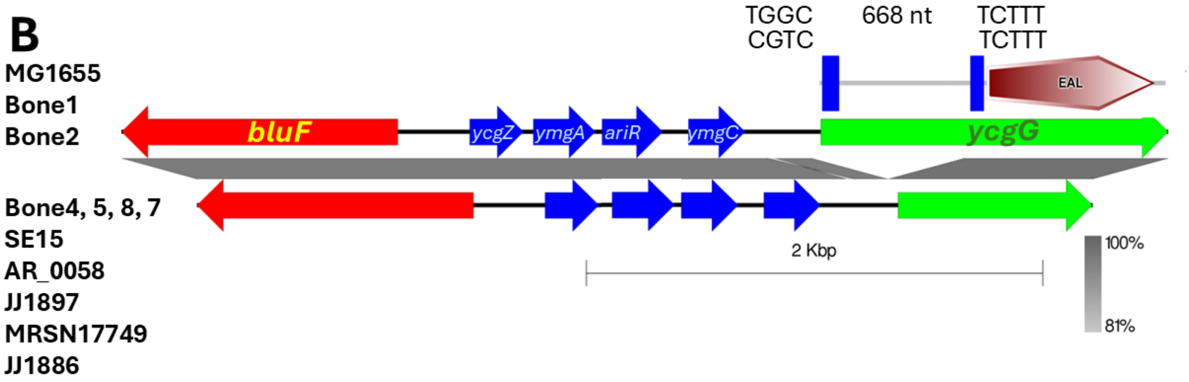

B Genomic context of the deletion of 668 bps resulting in the deletion of the CSS signaling domain of the phosphodiesterase YcgG creating a cytoplasmic only EAL protein.

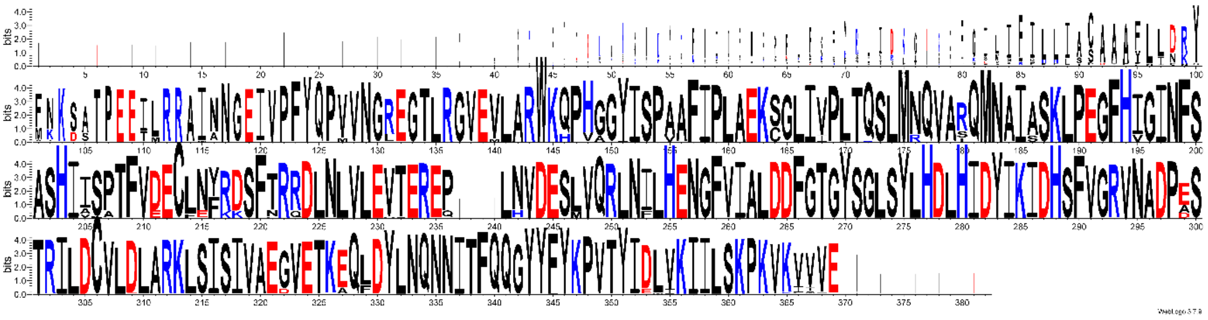

C WebLogo of aligned YcgG proteins from ST131 isolates retrieved by Blast search restricting the query to the ST131 database.

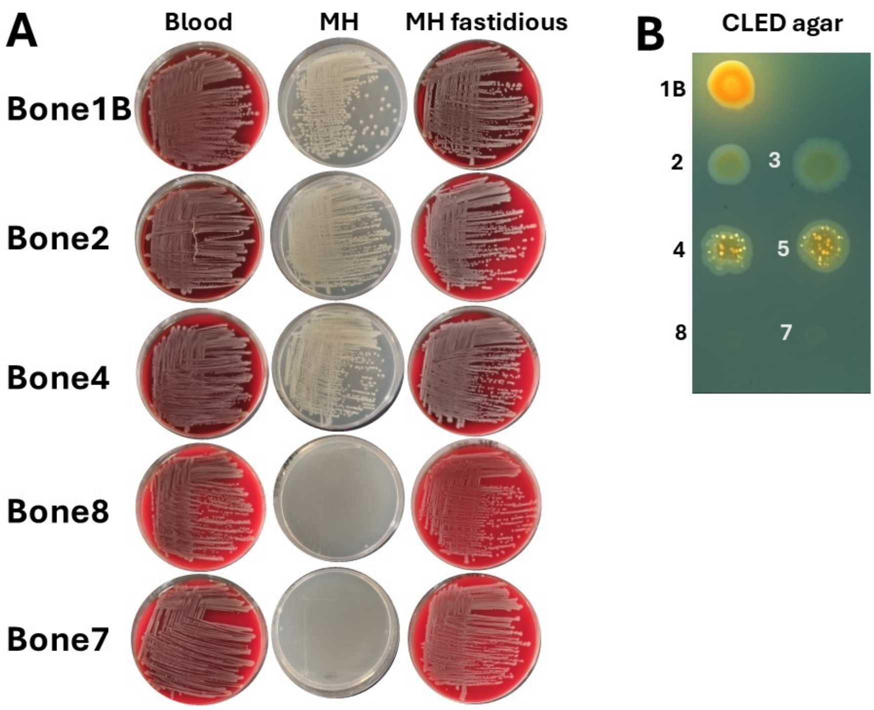

**Supplementary FIG 8** Growth characteristics of Bone isolates on agar media.
A Growth of Bone isolates on different agar media (left to right), blood agar, Mueller-Hinton agar and Mueller-Hinton fastidious agar after 24 h at 37 ºC. Bone8 and Bone7 isolates show a small colony variant phenotype on Mueller-Hinton agar plates.
B Growth of Bone isolates on CLED agar.

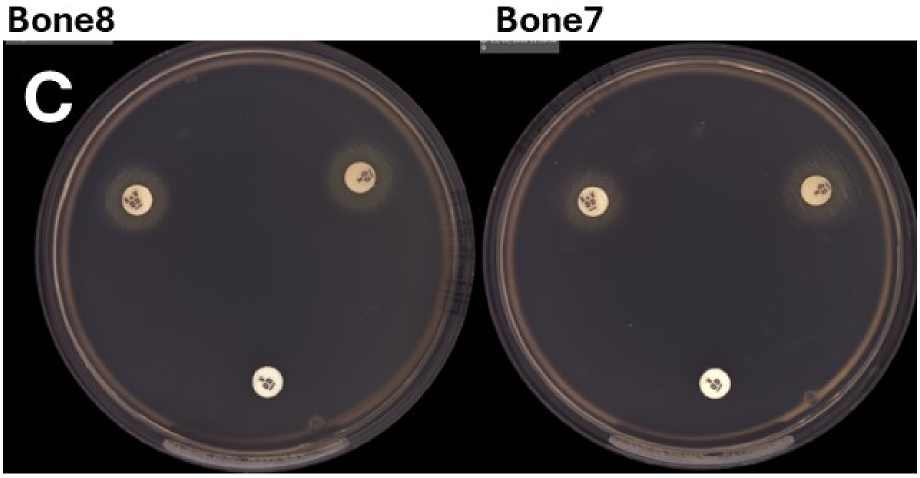

C Partial growth recovery of the small variant phenotype with hemin (factor X) and nicotinamide-adenine-dinucleotide (NAD; factor V) on Mueller Hinton broth after 24 h at 35º C.

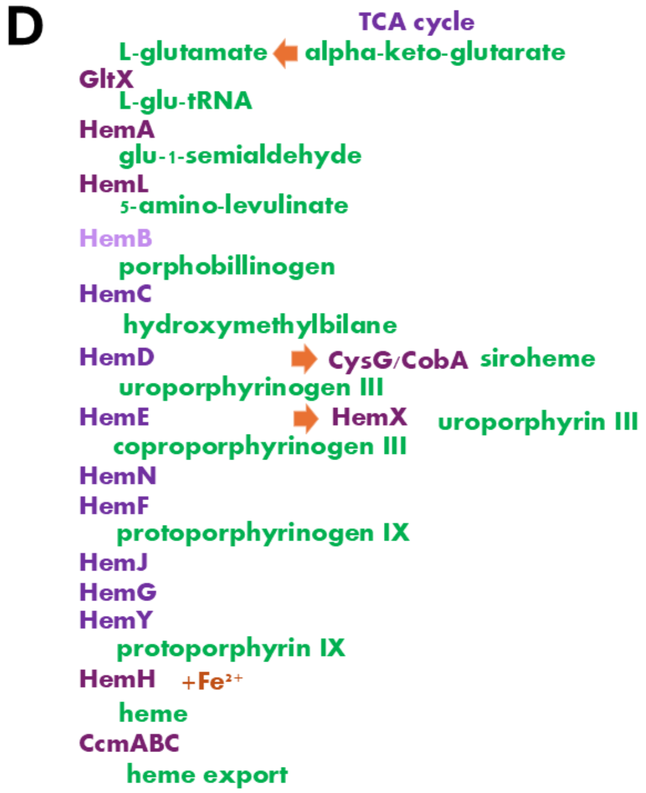

D Biosynthesis pathway leading to the synthesis of heme and siroheme. Impairment of HemB, the delta-aminolevulinic acid dehydratase not only leads to the lack of the prosthetic group heme.

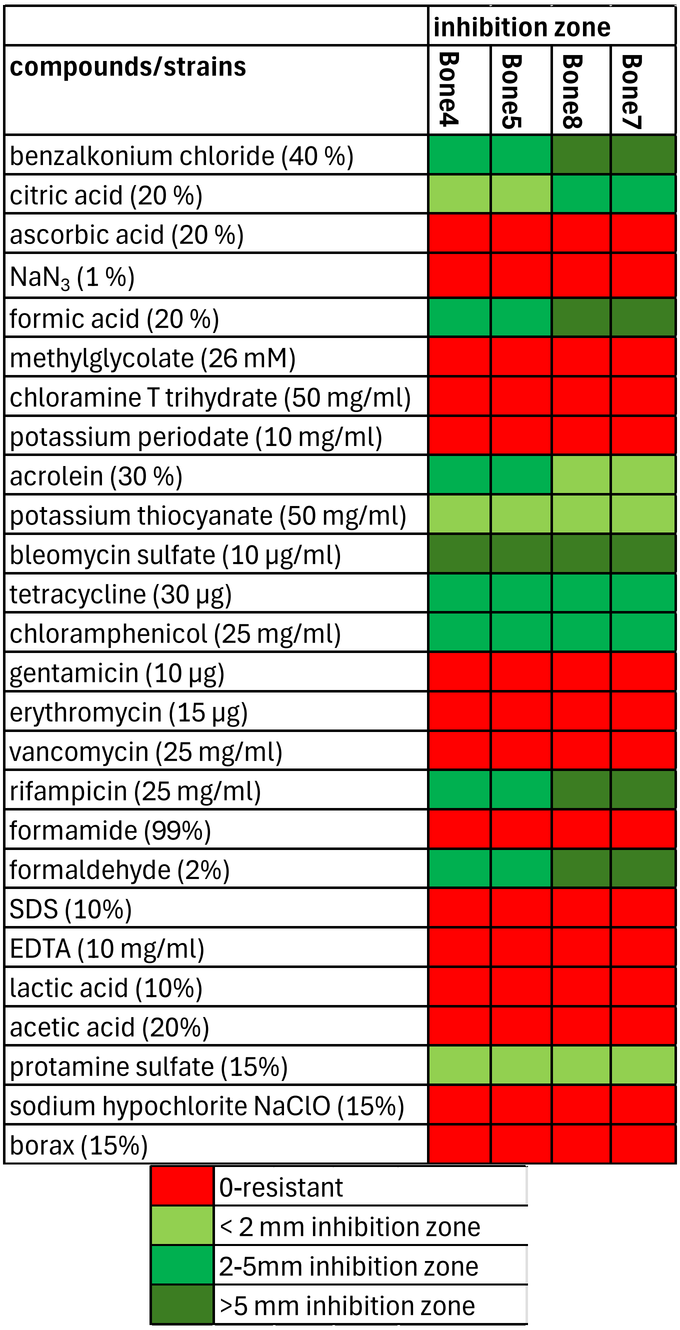

**Supplementary FIG 9** Susceptibility assay of ST131 Bone isolates for compounds relevant in tissue infection and antimicrobial treatment strategies.

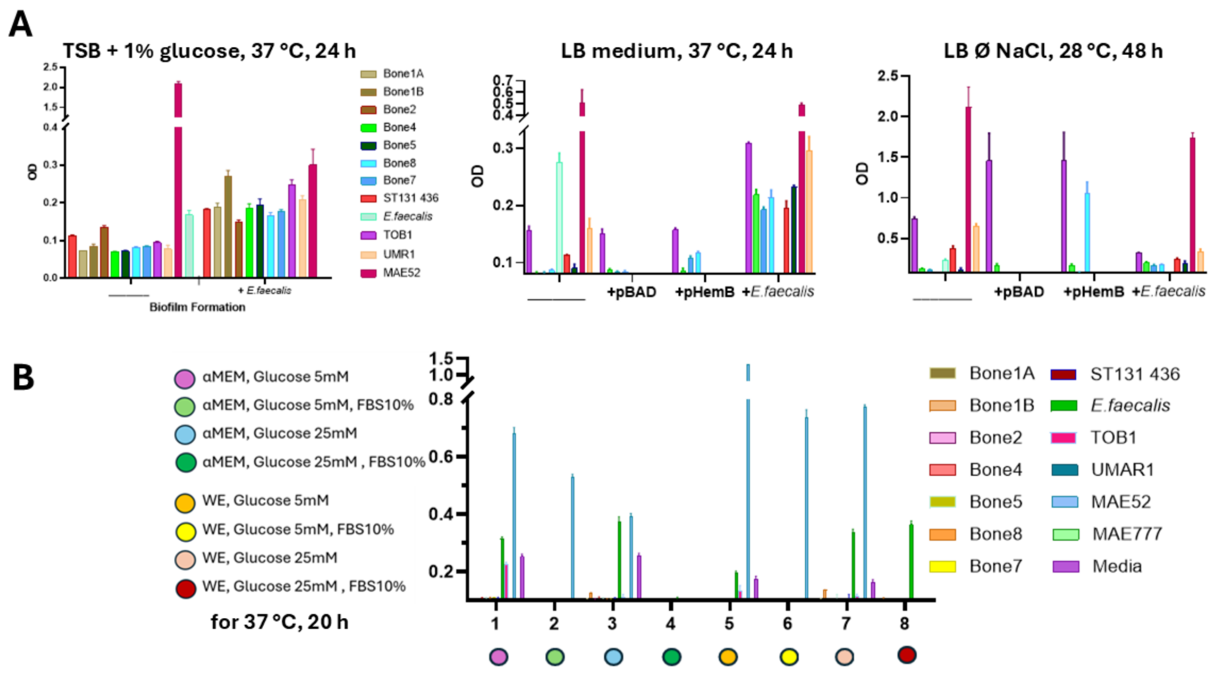

**Supplementary FIG 10** Biofilm formation and flagella-based swimming motility of Bone isolates in liquid medium.

A Biofilm formation of Bone isolates alone, complemented with pHemBCm or in coincubation with *E. faecalis* on a polystyrene surface in the 96-well plate assay.

B Biofilm formation on a polystrene surface in the 96-well plate assay upon incubation in cell culture medium. αMEM=complete minimum essential medium Eagle's α modification; WE=William's medium E. A and B, reference strains were ST131 EP436 (blood isolate), commensal *E. coli* TOB1 (phylogroup B 2), *S. typhimurium* UMR1, MAE52 (constitutive rdar biofilm) and MAE777 (no rdar biofilm).

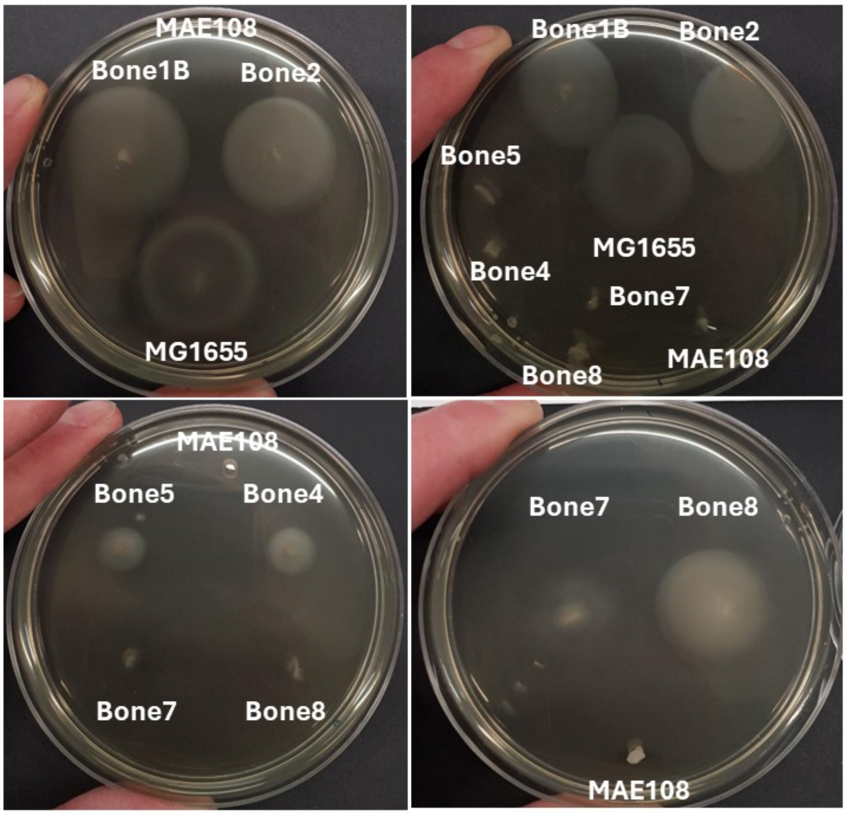

C Swimming motility in 0.25% Bolton broth base agar. Swimming motility of Bone1B, Bone2 and positive control MG1655 after 6.5 h, Bone4, Bone8, Bone7 and MG1655 after 9 h and Bone8 and Bone7 after 21 h of incubation at 37 ºC. MAE108=negative control.

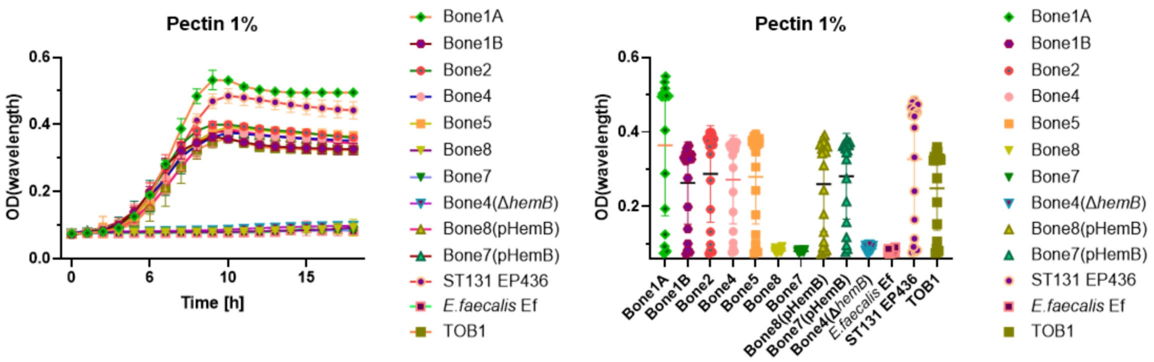

**Supplementary FIG 11** Assessment of pectin as carbon source for *E. coli* ST131 isolates. Bone isolates, Bone4,5,8 and 7 isolates complemented with pHemBCmBone, 4 Δ*hemB and E. faecalis* Ef were grown in M9 medium with 1% pectin (Kitchen) as sole carbon source for 24 h at 37 ºC. Reference strains were *E. coli* TOB1 and ST131 EP436. Left, time dependent growth curve; right display of maximal growth.
